# *Mycobacterium tuberculosis* resides in lysosome-poor monocyte-derived lung cells during chronic infection

**DOI:** 10.1101/2023.01.19.524758

**Authors:** Weihao Zheng, I-Chang Chang, Jason Limberis, Jonathan M. Budzik, B. Shoshana Zha, Zach Howard, Lucas Chen, Joel D. Ernst

## Abstract

*Mycobacterium tuberculosis* (Mtb) infects cells in multiple lung myeloid cell subsets and causes chronic infection despite innate and adaptive immune responses. However, the mechanisms allowing Mtb to evade elimination are not fully understood. Here, using new methods, we determined that after T cell responses have developed, CD11c^lo^ monocyte-derived lung cells termed MNC1 (mononuclear cell subset 1), harbor more live Mtb compared to alveolar macrophages (AM), neutrophils, and less permissive CD11c^hi^ MNC2. Bulk RNA sequencing of sorted cells revealed that the lysosome biogenesis pathway is underexpressed in MNC1. Functional assays confirmed that Mtb-permissive MNC1 have less lysosome content, acidification, and proteolytic activity than AM, and less nuclear TFEB, a master regulator of lysosome biogenesis. Mtb infection does not drive lysosome deficiency in MNC1 in vivo. Instead, Mtb recruits MNC1 and MNC2 to the lungs for its spread from AM to these cell subsets as a virulence mechanism that requires the Mtb ESX-1 secretion system. The c-Abl tyrosine kinase inhibitor nilotinib activates TFEB and enhances lysosome function of primary macrophages in vitro and MNC1 and MNC2 in vivo, improving control of Mtb infection. Our results indicate that Mtb exploits lysosome-poor monocyte-derived cells for in vivo persistence, suggesting a potential target for host-directed tuberculosis therapy.

**One Sentence Summary:** Virulent Mtb recruits and exploits intrinsically lysosome-deficient lung mononuclear cell subsets to resist elimination during chronic infection.

## INTRODUCTION

A distinguishing characteristic of *Mycobacterium tuberculosis* (Mtb) is its ability to evade elimination by innate and adaptive immune responses, leading to chronic infection with lung granuloma formation as a hallmark (*1*). Mtb’s ability to avoid elimination poses challenges to vaccine development (*1, 2*). Its persistence within host cells of lung granulomas leads to drug tolerance and suboptimal drug penetration(*3*), necessitating prolonged drug treatment for durable cure (*4–6*). Persistence and chronic infection also enable the prolonged transmission of Mtb and contribute to the ongoing global pandemic of tuberculosis (TB) (*7*). For these reasons, it is important to identify the permissive cellular niche of Mtb in vivo and determine the mechanisms that allow Mtb to persist in the face of innate and adaptive immunity.

Mtb is a facultative intracellular pathogen, and resides predominantly in mononuclear phagocytes, including resident tissue (i.e., alveolar) macrophages and in cells derived from circulating monocytes (*8–16*). There is also substantial evidence that the fate of pathogenic mycobacteria in distinct cell types can differ in vivo during chronic infection. Nearly 100 years ago, Florence Sabin reported two distinct cell types in vivo that differed in their handling of pathogenic mycobacteria: ‘clasmatocytes’ (tissue resident macrophages) “…phagocytize tubercle bacilli freely and fragment them”, while monocytes “retain the tubercle bacilli intact, with power to survive and multiply, over long periods of time” (*15*). These findings indicate that distinct types of mononuclear cells differ in their capacity to control pathogenic mycobacteria, but the identity of the cells that harbor Mtb and the mechanisms that determine their differential abilities to control Mtb during chronic infection are incompletely understood.

Recent studies using Mtb strains that constitutively express fluorescent proteins have confirmed that alveolar macrophages (AM), the tissue-resident macrophages of the air spaces, are the initial targets of infection (*9, 11, 14, 17*). During the initial 7-14 days of infection, Mtb replicates efficiently in AM in vivo (*9, 11, 12, 14*). However, the AM population is finite and does not expand markedly in response to infection (*12, 13, 16*). Therefore, for Mtb to expand its population and maximize the likelihood of transmission, the bacteria spread beyond AM.

One of the responses to Mtb infection is the recruitment of inflammatory cells, especially monocytes and neutrophils, to the lungs (*12, 13, 16, 18, 19*). Monocytes develop in the bone marrow (*20*) before entering the bloodstream, a step that depends on the chemokine receptor, CCR2 (*21*). In mice infected with Mtb, monocytes migrate from the blood to the lung parenchyma and differentiate into two distinct cell subsets distinguished by their expression of CD11c (*13*). CD11c^hi^ monocyte-derived cells were formerly considered dendritic cells (*16, 19, 22*), although they have also been considered closely related to macrophages based on their transcriptional profiles (*12*), while CD11c^lo^ monocyte-derived cells have been termed recruited macrophages (*16, 19, 23*). Both of these cell subsets become infected with Mtb within 3-5 days after they enter the lung parenchyma (*13*), and cells in both of these subsets increase in number and frequency in the lungs for at least 16 weeks post infection (*13*).

One consequence of the spread of Mtb from AM to other cell types is the transport of live bacteria from the lungs to the local draining lymph nodes (*22*) where bacterial antigens are transferred from infected migratory dendritic cells to uninfected resident lymph node dendritic cells for antigen-specific T cell priming (*24, 25*). Upon arrival of CD4 effector T cells in the lungs, the Mtb population stabilizes but is not eliminated. Together, these observations suggest that the cells in which Mtb resides after the acute stage of infection (≥4 weeks) cannot kill the bacteria at a rate greater than their growth, despite the presence of effector T cells.

Considering the events described above, the course of the host response to Mtb infection can be considered in at least 3 distinct stages. In mice, the initial stage is the first 10-14 days of infection, before developing antigen-specific CD4 or CD8 T cell responses. During the latter portion of the initial stage, inflammatory cells including neutrophils and monocytes are recruited to the lungs (*16, 19*). The second stage is transitional, comprising approximately 15-25 days post infection (dpi), and is marked by an accumulation of monocytes and their differentiation in the lung parenchyma, together with the appearance of effector CD4 and CD8 T cells. The third, chronic stage of infection, begins approximately 25-28 dpi, and is marked by further recruitment of effector T cells and a plateau in the number of bacteria in the lungs.

We developed and used new approaches to specifically quantitate viable Mtb in different lung myeloid subsets in the early chronic stage of infection (28 dpi). We found that CD11c^lo^ monocyte-derived cells we term MNC1 (mononuclear cell subset 1) harbor 4 to 6-fold more live bacteria per infected cell than other lung cell subsets. Although AM are an early replication niche for Mtb, they can kill Mtb after the initial stage of infection. We used RNA sequencing (RNA-seq) to identify differentially expressed genes and pathways that distinguish highly permissive MNC1 cells from less permissive CD11c^hi^ MNC2, restrictive AM, and neutrophils. This revealed that MNC1 express lower levels of lysosome biogenesis genes compared to AM, a finding we confirmed at the protein and functional levels. Furthermore, activation of the transcription factor TFEB by the c-Abl tyrosine kinase inhibitor nilotinib increases lysosome contents and functions of BMDM in vitro and MNC1 and MNC2 in vivo, improving control of Mtb. Together, our findings suggest that Mtb recruits and exploits lysosome-poor cells for persistence, and enhancement of lysosome function may be an effective strategy to improve the Mtb killing ability of permissive lung cells.

## RESULTS

### Characterization of lung cell populations containing Mtb after aerosol infection

We previously used flow cytometry to identify and characterize the lung leukocyte subsets containing fluorescent protein-expressing Mtb and found that at ≥ 14 dpi, the bacteria were found predominantly in neutrophils and in two subsets of monocyte-derived cells which we termed ‘recruited macrophages’ and ‘myeloid dendritic cells’ (*16*). Since subsequent studies have identified markers that allow higher resolution definition of lung leukocytes (*13*), we repeated those studies, using flow cytometry to analyze lung cells from mice infected with aerosol Mtb H37Rv expressing ZsGreen at 14, 21, and 28 dpi (Fig. 1A-1C and Fig. S1). The results were consistent with reports that the number of AM changed minimally over the 28 days of infection, while the number of neutrophils, MNC1, and MNC2 in the lungs markedly increased by 21-28 dpi (Fig. 1A). As we previously reported (*16, 26*), the number of infected (ZsGreen^+^) neutrophils and MNC2 exhibited a transient peak 21 dpi, followed by a decrease by 28 dpi. In contrast, the number of infected MNC1 cells increased from 14 dpi to 28 dpi (Fig. 1B). When considered as the fraction of the total number of Mtb-infected cells, AM were dominant 14 dpi consistent with prior results (*9, 11, 14*). By 21 dpi, AM constituted a minor fraction of the total, and neutrophils, MNC1, and MNC2 were the predominant populations that contained Mtb (Fig. 1C). At this time point, MNC2 were the dominant subset of infected cells (Fig. 1C), consistent with a recent report that used sfYFP-expressing Mtb and a similar flow cytometry scheme (*12*). However, MNC1 expanded further as a fraction of the infected cells at 28 dpi. It is also notable that regardless of the cell subset, only a minority of the cells in a subset contain Mtb. Overall, the data confirm that AM are the primary Mtb-infected population in the initial stage of aerosol infection, and that MNC1 and MNC2 are the major infected cell subsets after the initial stage.

**Fig 1.**
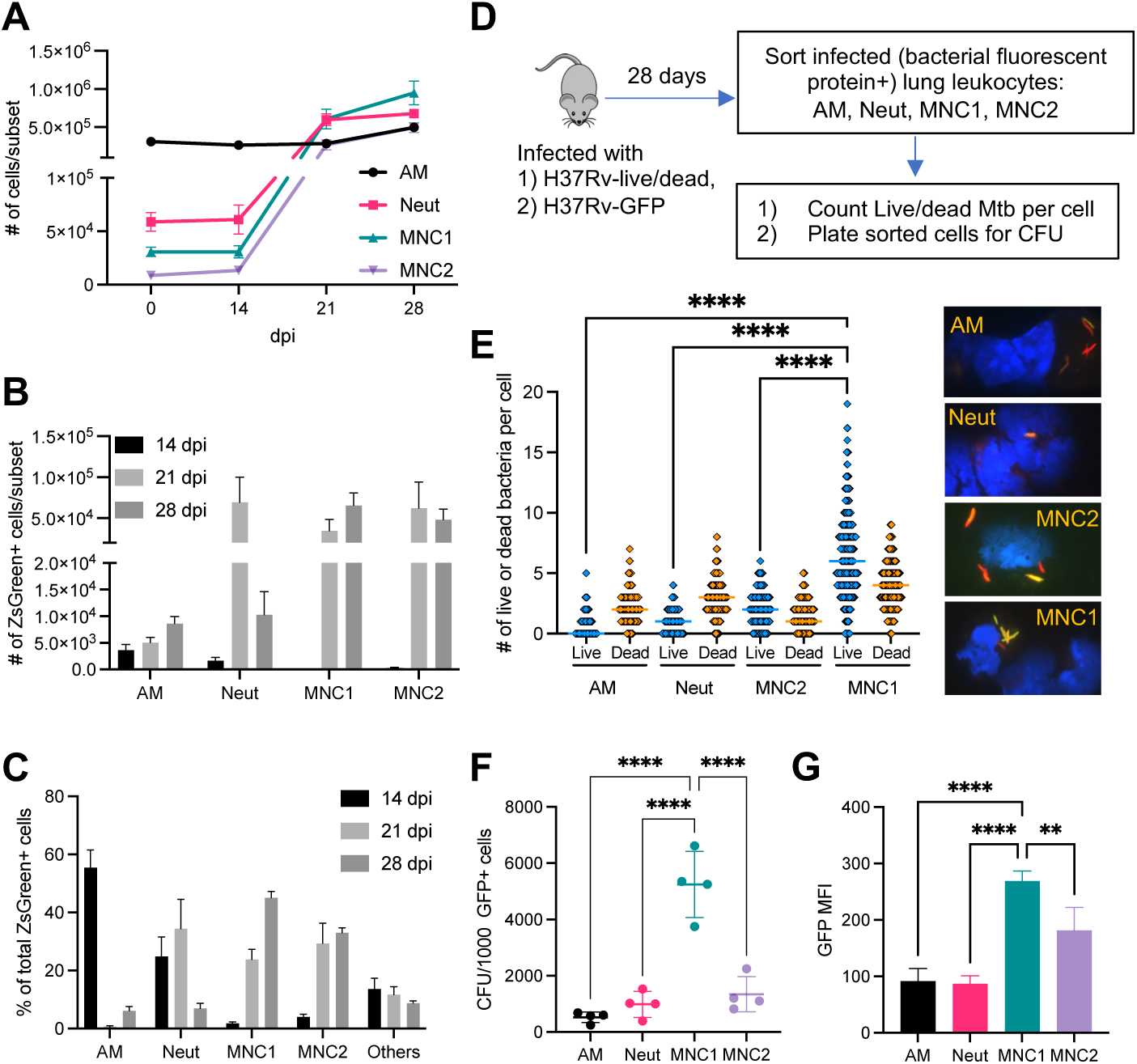
MNC1 are highly permissive for Mtb intracellular survival. C57BL/6 mice were infected with the designated strain of Mtb by low-dose aerosol. Lung cells were isolated for flow cytometry analysis or live cell sorting at the indicated timepoints post infection. **(A)** Lung phagocyte population dynamics after Mtb (H37Rv-ZsGreen) infection. Neutrophils (Neut), MNC1, and MNC2 increase in response to Mtb infection, while the number of alveolar macrophages (AM) changed minimally. **(B)** Number of Mtb H37Rv-ZsGreen^+^ cells in distinct lung phagocyte subsets by flow cytometry. **(C)** Cell type composition of total Mtb H37Rv-ZsGreen^+^ lung leukocytes by flow cytometry. After predominant distribution of Mtb in AM and neutrophils, MNC1 and MNC2 dominate by 28 dpi. **(D)** Schematic diagram of procedures to quantitate intracellular live Mtb in sorted lung phagocyte subsets. C57BL/6 mice were infected with Mtb H37Rv-live/dead or H37Rv-GFP and cells containing fluorescent protein-expressing bacteria were analyzed at 28 dpi. H37Rv-live/dead carries a plasmid that drives constitutively expression of mCherry and tetracycline-inducible GFP. For induction of GFP expression in live bacteria, doxycycline was administered via drinking water for 6 days before harvest at 28 dpi. **(E)** MNC1 contain the largest number of live Mtb per cell 28 dpi. Quantitation of live (GFP^+^mCherry^+^) or dead (GFP^-^mCherry^+^) Mtb per infected cell (n ≥ 300) was performed by fluorescence microscopy on viable cells sorted from mice infected with Mtb H37Rv-live/dead. Representative images on the right show live and dead Mtb. Dead (mCherry^+^GFP^-^) Mtb are red; live (mCherry^+^GFP^+^) appear yellow. The majority of the Mtb in AM and neutrophils are dead, while the majority of the bacteria in MNC1 and MNC2 are live (MNC1>MNC2). **(F)** MNC1 contain the largest number of live Mtb (H37Rv-GFP) at 28 dpi. Cells in each subset were sorted according to surface phenotypes and for bacterial status (GFP^+^). CFU of sorted GFP^+^ cells in each subset were counted after 3 wk incubation. The results are expressed as CFU per 1000 GFP^+^ cells in each subset. **(G)** GFP MFI correlates with live Mtb burdens in the 4 infected lung myeloid cell subsets from mice infected with H37Rv-GFP (28 dpi). Unless otherwise stated (e.g., MFI), results are presented as mean ± SD of 4-5 mice, and are representative of 2-3 independent experiments. **p<0.01 ****p<0.0001 by one-way ANOVA.

MerTK and CD64 have been used as markers to define CD11b^lo^ AM and CD11b^+^ lung interstitial macrophages (IM) in various contexts, including mice intranasally infected with a high dose of Mtb (*11, 27–29*). Since strong evidence indicates that MNC1 and MNC2 are derived from monocytes (*12, 13, 19*), we used a modified flow panel (Fig. S2A) to query if MNC1 and MNC2 subsets are similar to IM defined using MerTK and CD64 expression. This revealed that ∼95% of AM were MerTK^+^CD64^+^, while only 10-22% of MNC1 and 20-47% of MNC2 were MerTK^+^CD64^+^ from 14-28 dpi (Fig. S2B). When we gated on Mtb^+^ cells, we found that 70-97% of infected AM were MerTK^+^CD64^+^, while fewer MNC1 (10%-60%) and MNC2 (25%-86%) were MerTK^+^CD64^+^. These results indicate that gating out cells that are not MerTK^+^CD64^+^ excludes substantial fractions of the Mtb-infected cells in the lungs.

### Cell subset distribution of live Mtb during the chronic stage of infection

Use of fluorescent protein-expressing strains of Mtb coupled with flow cytometry has been invaluable in revealing the diversity of the cell types that are infected in vivo, but these procedures alone do not reveal the viability of the intracellular bacteria. Likewise, although there is evidence that cells exhibiting brighter bacterial fluorescence harbor more bacteria (*12*), a given cell can contain both live and dead Mtb that contribute to the fluorescence signal. To quantitate live intracellular Mtb, we utilized Mtb H37Rv carrying a live/dead reporter plasmid that drives constitutive expression of mCherry fluorescent protein (all bacteria) and doxycycline-inducible expression of green fluorescent protein (GFP; only live bacteria) (*30*). Using fluorescent cell sorting and fluorescence microscopic evaluation, we enumerated live (mCherry^+^GFP^+^) and dead (mCherry^+^GFP^-^) bacteria in individual infected cells (Fig. 1D).

We sorted four lung cell subsets from mice infected with live/dead-H37Rv (28 dpi) and examined mCherry^+^ Mtb in flow-sorted AM, neutrophils, MNC1, and MNC2 by fluorescence microscopy. This revealed that the majority of the bacteria in mCherry^+^ AM were dead (GFP^-^) at this time point (Fig. 1E). Within the mCherry^+^ AM population, there was considerable variation in the number of total bacteria (range: 1-7) per cell. While most of the mCherry^+^ AM contained both live and dead bacteria, some contained only dead bacteria and others contained only live bacteria. Overall, dead bacteria were more abundant than live bacteria in AM. These findings are consistent with the recent report that vaccination with intrapulmonary BCG causes the arrest of growth and restricts the spread of virulent Mtb from AM (*31*). They are also consistent with the results indicating that Mtb expansion in AM appears to be arrested by 21 dpi (*12*). Similar results were apparent in the neutrophil population, in which there was also a range of bacteria per mCherry^+^ cell; most contained both live and dead bacteria, with dead bacteria predominating.

In contrast to the bacterial states in AM and neutrophils, in both monocyte-derived cell subsets, MNC1 and MNC2, live bacteria were more abundant than dead bacteria (Fig. 1E). Some MNC2 cells contained single live or dead bacteria, while most contained multiple bacteria, including both live and dead bacteria in the same cell. Since the MNC2 cell subset resembles the CD11c^hi^ cell subset previously reported to represent the largest fraction of YFP-expressing Mtb at 21 dpi (*12*), these results confirm that this (or a related) subset contains predominantly live bacteria at a later time point (28 dpi). The distribution of Mtb in the MNC1 subset was similar to that in the MNC2 subset, although MNC1 cells contained more live bacteria per mCherry^+^ cell. Very few infected MNC1 cells contained single bacteria, while the median number of either live or dead bacteria per mCherry^+^ cell exceeded that observed in any of the other cell subsets, including MNC2. Notably, the median number of live Mtb per mCherry^+^ MNC1 cell (6; range, 0-19) was higher than that of the dead bacteria (4; range, 0-9), indicating that, although each of the sorted lung cell subsets exhibited the ability to kill some virulent Mtb, MNC1 cells are the least restrictive for intracellular growth of Mtb at a time point (28 dpi) when T cell responses are well developed and the total size of the bacterial population in the lungs has stabilized (*22*).

To extend these results, we performed a similar experiment using Mtb H37Rv constitutively expressing enhanced GFP (*16*), and quantitated live Mtb as colony-forming units (CFU) present in sorted GFP^+^ cells in each of the subsets (Fig. 1D). This revealed that 28 dpi, AM and neutrophils both contained ∼400-600 CFU/1,000 GFP^+^ cells, MNC2 contained approximately 1,000 CFU/1,000 GFP^+^ cells, and MNC1 contained 4,000-6,000 CFU/1,000 GFP^+^ cells (Fig. 1F). The GFP or ZsGreen median fluorescence intensity (MFI) of infected MNC1 was higher than the other infected subsets, correlating with CFU and the number of live Mtb per cell in each infected subset (Fig. 1G and Fig. S3A). Moreover, at 56 dpi, Mtb resided predominantly in MNC1, which also harbored more Mtb per cell than other subsets as indicated by bacterial fluorescence (Fig. S3B-S3C).

Together, these results suggest that during the chronic stage of infection, after the development of adaptive T cell responses, AM and neutrophils restrict and kill virulent Mtb effectively, although they do harbor some live bacteria. In contrast, MNC1 and MNC2 are permissive for Mtb, as they predominantly harbor live bacteria and MNC1 are more permissive than MNC2 cells.

### RNA-seq analysis of lung myeloid subsets reveals diversity and differential gene and pathway expression

To identify mechanisms that potentially account for the differential ability of lung cell subsets to restrict and kill Mtb during chronic infection, we carried out RNA-seq analysis for 8 cell populations sorted from the lungs of Mtb-infected mice: AM, neutrophils, MNC1, and MNC2; each either infected or bystander (uninfected). Although we found differentially expressed genes between infected and bystander cells for each subset (Fig. S4), a t-stochastic neighbor embedded (tSNE) plot of the RNA-seq data showed segregation of the four cell subsets (Fig. 2A). In this analysis, Mtb-infected cells were subclusters within each cell subset, indicating that the presence of intracellular bacteria does not determine the cell phenotype. The transcriptome data revealed that canonical markers *Ear1*, *Mrc1*, *Pparg*, *Siglecf*, *Siglec1*, and *Fabp4* were indeed highly expressed by AM, and neutrophils expressed *Il1r2*, *Csf3r*, *Cxcr2*, *S100a8*, *S100a9* and *Retnlg* (Fig. 2B). In contrast, monocyte markers such as *Ccr2*, *Cx3cr1*, *Mafb*, *Ly6c1* and *Tfrc* were expressed at a significantly higher level in MNC1 and MNC2 compared to AM, consistent with other evidence that MNC cells are derived from monocytes (*13*). MNC2 also expressed transcripts characteristic of DC including *Ccl22*, *Il12b*, *Flt3*, *Ccr7*, *Cd83*, and *Cd86*. However, other studies have indicated that classical DC are much less abundant than monocyte-derived cells in the lungs of Mtb-infected mice (*27*), and that CD11c^hi^ monocyte-derived cells (MNC2 in this study) also express transcripts characteristic of macrophages (*12*), indicating that this population may be heterogenous.

**Fig. 2.**
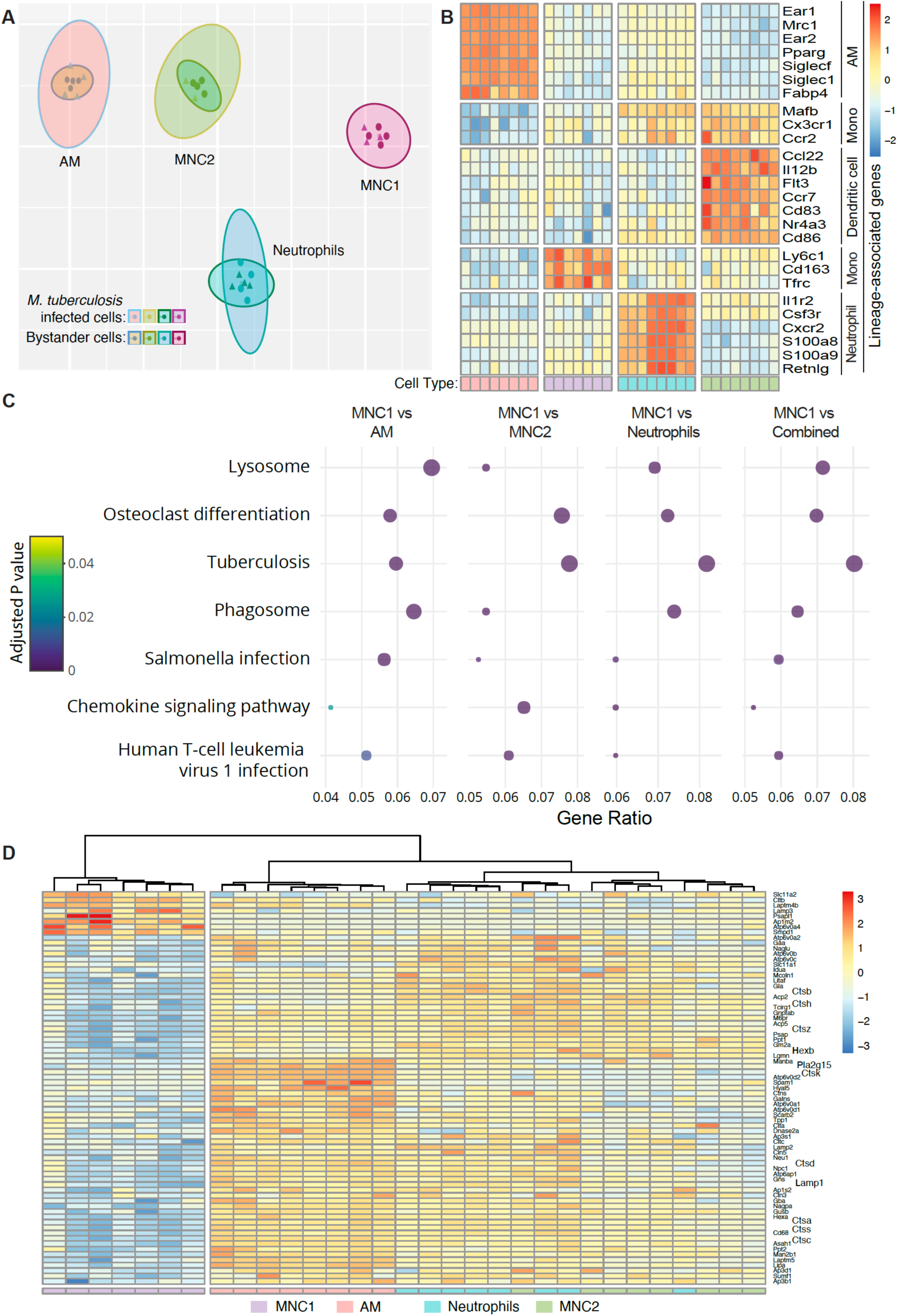
RNA-seq analysis of live-sorted phagocyte subsets from lungs of Mtb-infected mice reveals evidence of deficient lysosome biogenesis in MNC1. C57BL/6 mice were infected with Mtb H37Rv-mCherry by low dose aerosol. After 28 days, 4 pools of lung cells from 5 infected mice per pool were prepared and stained, and 10,000 live cells from each subset in each of the 4 pools were sorted directly into RNA later and processed for bulk RNA sequencing. **(A)** t-stochastic neighbor embedding (t-sne) plot showing distinct clusters of four myeloid cell types based on RNA sequencing on cells sorted from lungs of Mtb H37Rv-mCherry infected mice. Within a clustered subset there is substantial overlap between Mtb infected and bystander cells after exclusion of one infected MNC1 outlier sample. **(B)** Heatmap showing separation of four distinct cell types based on Z-scores from variance stabilized read counts for lineage markers. The color coding of the cell types shown at the bottom correspond to the colors in (A). **(C)** Dot plot showing 7 of 18 KEGG pathways that differ significantly with an enrichment ratio greater than 0.04 for AM, MNC2, neutrophils, combined analysis (AM, MNC2, neutrophils) and MNC1. The color represents the adjusted p values, the graph is ordered by descending values (lowest p value = 1.5 × 10^-16^ for the lysosome pathway, combined analysis), while the dot size is proportional to the gene count. **(D)** Heatmap of the Z-scores from variance stabilized read counts for significantly differentially expressed genes of KEGG lysosome pathway shows separation of MNC1 from AM, MNC2, and neutrophils.

We performed KEGG pathway analysis to identify differences contributing to the differential Mtb permissiveness of MNC1 compared with the other 3 lung cell subsets. We identified the pathways that exhibited >2-fold enrichment with a Benjamini-Hochberg adjusted p<0.05 (Fig. 2C and Fig. S5). Of these, the “Lysosome” pathway was differentially expressed in MNC1 compared with the other subsets (Fig. 2C). Genes of lysosome pathways encode lysosomal membrane proteins (e.g., LAMP1, LAMP2), lysosomal hydrolases (e.g., cathepsin proteases and glycosidases), and lysosome vacuolar proton ATPase (V-ATPase) subunits (Fig. 2D). These genes were not differentially expressed in Mtb-infected cells compared to bystander cells within each subset, and 65 of 73 genes were underexpressed in both Mtb-infected and bystander MNC1 cells. Among the underexpressed lysosome genes, beta-hexosaminidase (HEXB), cathepsins B, S, and L, and phospholipase A2 (PLA2G15) have been reported to contribute to antimycobacterial activity (*32–34*). Notably, some of the underexpressed lysosome genes such as V-ATPase subunits are also important components of the KEGG “Tuberculosis” and “Phagosome” pathways. Thus, we hypothesized that the intrinsic deficiency of lysosome biogenesis in MNC1 is a mechanism that contributes to their Mtb permissiveness.

### Mtb-permissive MNC1 are deficient in lysosomal enzyme activity

Since Mtb-restrictive AM and Mtb-permissive MNC1 cells exhibit the greatest difference in their expression of genes involved in lysosome biogenesis, and since Mtb survives in macrophages at least in part by limiting lysosome-dependent phagosome maturation (reviewed in (*2*)) and lysosome-dependent autophagy (*35–38*), we quantitated lysosome activities and content in AM and MNC1.

We first used a fluorogenic Cathepsin B (CTSB) assay and flow cytometry to compare the enzymatic activity of CTSB in AM and MNC1 from lungs of mice infected with Mtb for 28 days. In this assay, the substrate fluoresces after cleavage and can be quantitated by fluorescence microscopy or flow cytometry as a reflection of cathepsin B enzymatic activity (*39*). When incubated with the CTSB substrate, AM showed ∼7-fold higher MFI of product than MNC1 (Fig. 3A-3B). We then sorted AM and MNC1 from Mtb-infected mice, incubated them with the CTSB substrate, and quantitated fluorescence by microscopy. In this assay, AM incubated with the substrate exhibited red fluorescence that was readily detectable by fluorescence microscopy (Fig. 3C, left panels). In contrast, sorted MNC1 cells exhibited barely detectable fluorescence. As a control, the specific V-ATPase inhibitor bafilomycin A1 that prevents lysosomal acidification and lysosomal cathepsin activity (*40*), blocked generation of fluorescence in AM (Fig. 3C, right panels). Quantitative image analysis revealed ∼3-4-fold higher CTSB product fluorescence in AM compared with MNC1 (Fig. 3D).

**Fig 3.**
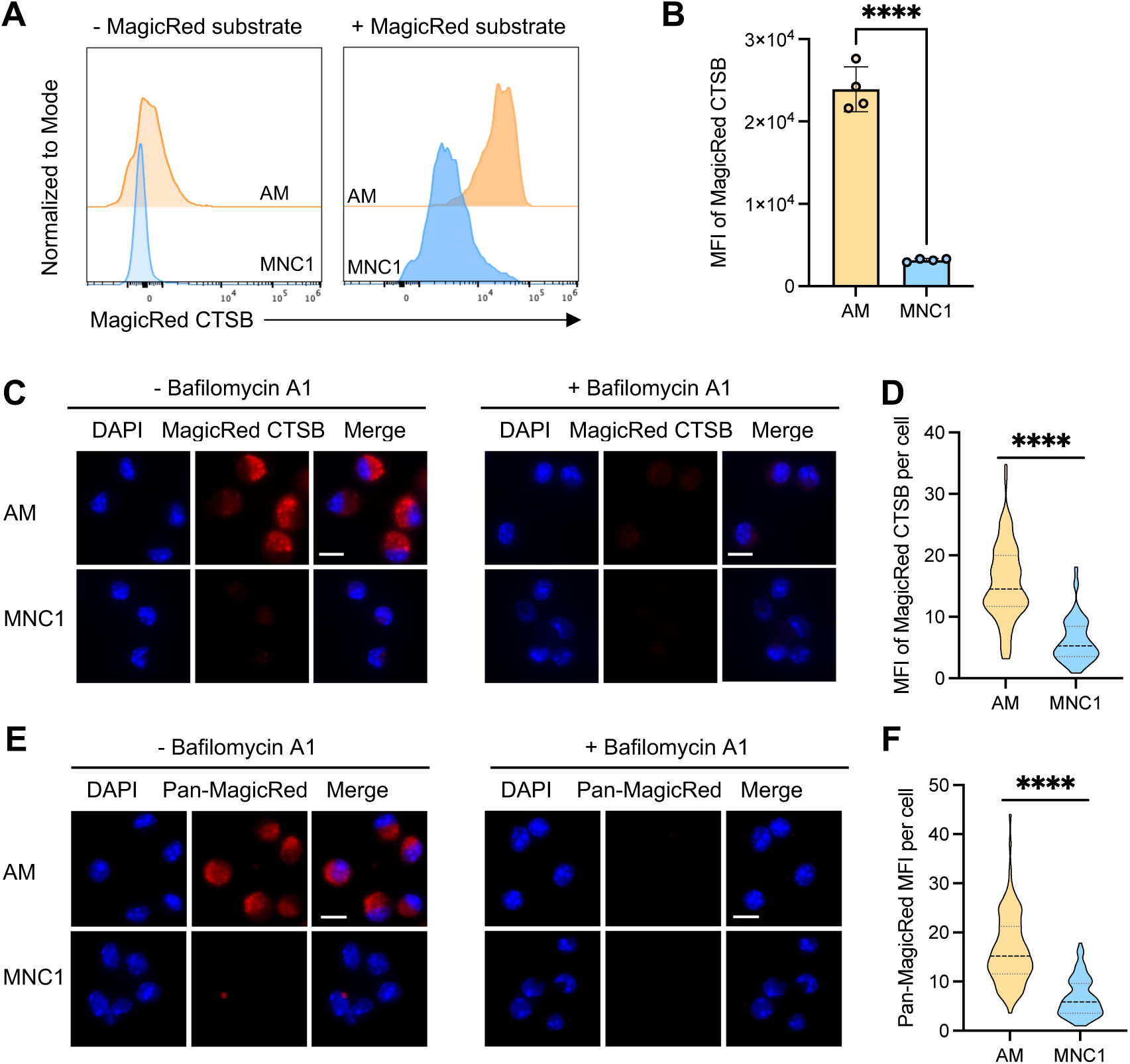
MNC1 cells are deficient in lysosomal cathepsin proteolytic activities. C57BL/*6* mice were infected by aerosol with ∼100 Mtb H37Rv-ZsGreen or H37Rv-mCherry. At 28 dpi, mouse lungs were harvested for flow cytometry analysis, live cell sorting, and fluorogenic cathepsin (MagicRed^®^) assays. (**A-B**) Representative histograms and MFI of the fluorescent product of CTSB activity for AM and MNC1. Lung cells were isolated from mice infected with H37Rv-ZsGreen (28 dpi), then incubated with MagicRed^®^ CTSB substrate for 30 min, and stained with antibodies for discrimination of cell subsets, followed by flow cytometry analysis. **(C)** CTSB activities in AM and MNC1 sorted from mice infected with H37Rv-mCherry for 28 days, analyzed by fluorescence microscopy. Live-sorted cells from each cell subset were treated with the fluorogenic CTSB substrate in the absence or presence of Bafilomycin A1 (BafA, 100 nM) for 1 h. Cells were fixed and analyzed by confocal microscopy. Scale bars, 10 μm. **(D)** Quantification of fluorogenic CTSB product MFI per cell from the left panel in (C). >100 cells per subset were analyzed using ImageJ. **(E)** Pan-cathepsin activities in AM and MNC1 sorted from lungs of mice infected with H37Rv-mCherry (28 dpi). Sorted cells were treated with a pool of MagicRed^®^ fluorogenic cathepsin substrates (CTSB+CTSK+CTSL) in the absence or presence of 100 nM BafA for 1 h. Cells were fixed and analyzed by fluorescence microscopy. Scale bars, 10 μm. **(F)** Quantification of Pan-cathepsin product MFI per cell from the left panel of (E). >100 cells per subset were analyzed using ImageJ. ***p<0.0001 by unpaired Student’s t test. Results are presented as mean ± SD, representative of 2-3 independent experiments.

Since RNA-seq analysis revealed decreased expression of mRNA encoding other lysosomal cathepsins (H, Z, K, D, A, S, L, and C) (Fig. 2D and Data file S1), we analyzed additional lysosomal cathepsin activities using a pool of substrates for cathepsins B, K, and L. This yielded results similar to those obtained with the CTSB substrate: fluorescent product generation was ∼3-fold greater in AM than in MNC1 cells, and the fluorescence generation in AM was abrogated by bafilomycin A1 (Fig. 3E and 3F). Together, these results provide functional evidence for the lower expression of lysosomal cathepsin mRNAs in MNC1 compared with AM.

### Mtb-permissive MNC1 are deficient in lysosomal acidification

Lysosomal hydrolases and antimicrobial activities (*41, 42*) require an acidic environment for their functions; the acidic lysosome environment is provided by the activity of a V-ATPase that comprises multiple V0 and V1 protein subunits (*43*). Although the mRNA level of V-ATPase subunits did not differ significantly different between infected and bystander cells, RNA-seq revealed lower expression of multiple V-ATPase subunits in MNC1 compared with AM (Fig. 4A). Thus, we hypothesized that MNC1 are deficient in lysosomal acidification, contributing to their deficient lysosome activity.

**Fig 4.**
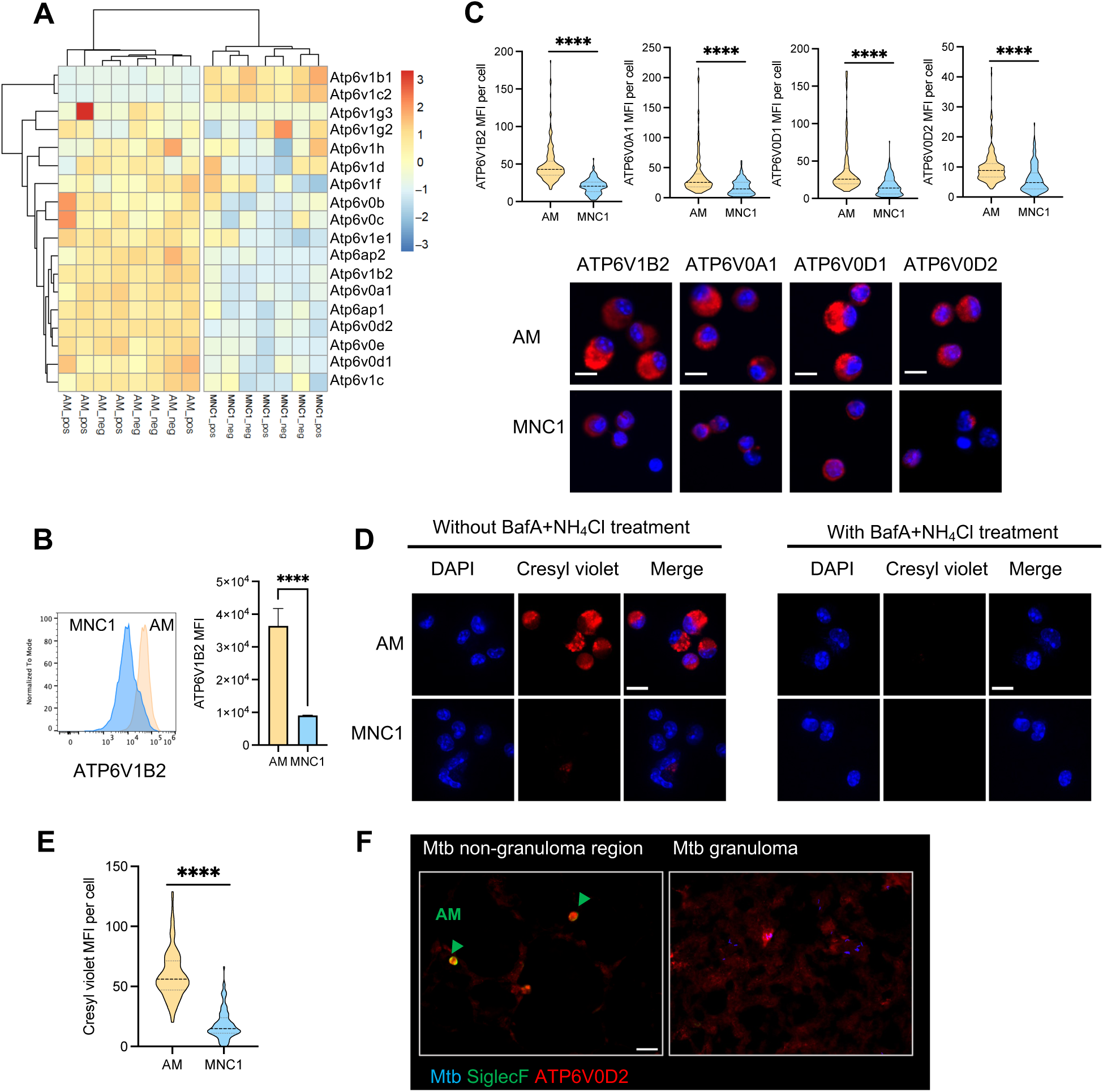
MNC1 cells exhibit defective lysosome acidification. C57BL/6 mice were infected by aerosol with ∼100 Mtb H37Rv-ZsGreen or H37Rv-mCherry. At 28 dpi, mouse lungs were harvested for cresyl violet staining and flow cytometry analysis or live cell sorting, and fluorescence microscopy. **(A)** Heatmap of the Z-scores from variance stabilized read counts for differentially expressed vacuolar proton ATPase (V-ATPase) subunit genes in AM vs MNC1. “_pos” indicates sorted cells containing Mtb; “_neg” indicates bystander cells that did not contain bacteria. **(B)** Representative histograms and MFI of ATP6V1B2 protein immunostaining by flow cytometry of fixed and permeabilized AM and MNC1 from mice infected with H37Rv-ZsGreen (28 dpi). **(C)** Immunofluorescence analysis of four V-ATPase subunits in AM and MNC1 sorted from H37Rv-mCherry-infected mice (28 dpi). Representative images were taken by confocal microscopy. Scale bars, 10 μm. Quantification of V-ATPase subunit MFI per cell from >100 cells per subset was done using ImageJ. **(D)** Cresyl violet assay of lysosome acidification in AM and MNC1 sorted from lungs of mice infected with H37Rv-mCherry (28 dpi). Sorted cells were treated with 5 µM cresyl violet in the absence or presence of BafA (100 nM) and NH_4_Cl (10 mM) for 30 min. Cells were fixed and analyzed by confocal microscopy. Scale bars, 10 μm. **(E)** Quantification of cresyl violet MFI per cell for the left panel in (D) using ImageJ (>100 cells of each type). **(F)** ATP6V0D2 and SiglecF staining of lung sections from mice infected with H37Rv-mCherry (28 dpi). Scale bar, 20 μm. Green arrow indicates SiglecF^+^ AM. ****p<0.0001 by unpaired Student’s t test. Data are presented as mean ± SD, representative of 2-3 independent experiments.

We first analyzed the protein level of V-ATPase subunits in AM and MNC1 isolated from the lungs of Mtb-infected mice (28 dpi). By flow cytometry, intracellular ATP6V1B2 was ∼3 fold higher in AM than in MNC1 (Fig. 4B). As assessed by immunofluorescence microscopy on sorted AM and MNC1, all of the V-ATPase subunits that we examined (ATP6V1B2, ATP6V0A1, ATP6V0D1, and ATP6V0D2) were present at 2-4 fold higher levels in AM compared with MNC1 (Fig. 4C). These results are consistent with the results of RNA analyses, indicating that MNC1 may be less capable of lysosome and phagolysosome acidification compared with AM.

To determine whether the lower abundance of V-ATPase subunits has functional significance for lysosome acidification, we quantitated accumulation of the anionic (pKa = 9.84) fluorochrome, cresyl violet, which labels acidic compartments (*44*). Fluorescence microscopy revealed marked accumulation of cresyl violet in punctate structures in AM, but minimal accumulation in MNC1 cells (Fig. 4D-4E). Treatment of AM with bafilomycin A1 and ammonium chloride to block lysosome acidification abrogated accumulation of cresyl violet, confirming that cresyl violet accumulation and fluorescence are the consequence of lysosome acidification mediated by V-ATPase activity.

We next used immunofluorescence to characterize expression of the V-ATPase subunit ATP6V0D2 in lung sections from Mtb-infected mice (28 dpi). Using CD11b and SiglecF antibodies, we first confirmed that AM were absent from granulomas, located primarily in non-granuloma lung regions, while the majority of granuloma cells were SiglecF^-^CD11b^+^ (Fig. S6A), coinciding with previous findings (*9, 13*). Since CD11b is also expressed on neutrophils, we used F4/80 and CD68 along with CD11b to stain the Mtb lung sections. F4/80 and CD68 colocalized with most of the CD11b^+^ cells including Mtb-infected cells (Fig. S6B), providing evidence that the SiglecF^-^CD11b^+^ cells are macrophages that likely represent monocyte-derived MNC1 and/or MNC2 cells. In these sections, AM exhibited a higher level of ATP6V0D2 than granuloma cells in the infected lungs (Fig. 4F). These results support the flow cytometry evidence that monocyte-derived cells in granulomas are deficient in lysosomal proteins.

### Mtb-permissive MNC1 are deficient in lysosome enzyme content and LAMP1+ organelles

Our finding that lysosomal cathepsin activity is reduced in MNC1 compared with AM could be secondary to reduced lysosome acidification, or due to decreased abundance of lysosomal enzyme protein. By immunofluorescence microscopy, we found abundant CTSB staining in a punctate distribution in AM, while CTSB-containing puncta were less numerous in MNC1, and this was substantiated by quantitative image analysis (Fig. 5A-5B). These findings indicate that deficient lysosome acidification alone is unlikely to account for the lower cathepsin activity in MNC1 compared with AM, and, consistent with lower mRNA expression, immunoreactive CTSB protein is also less abundant in MNC1 than in AM.

**Fig 5.**
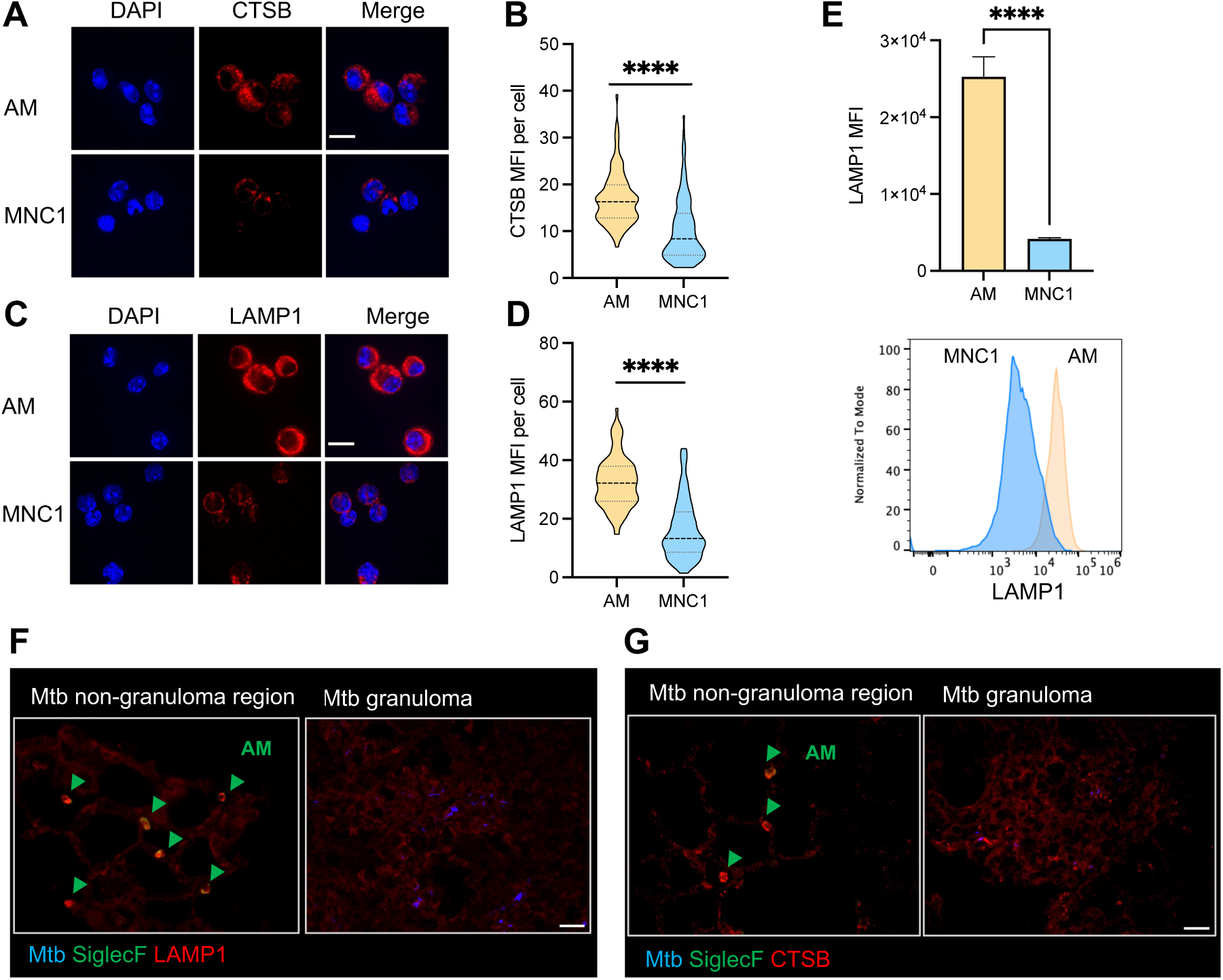
MNC1 are deficient in lysosomal cathepsin B and LAMP1 protein content. C57BL/6 mice were infected by aerosol with ∼100 Mtb H37Rv-ZsGreen or H37Rv-mCherry. At 28 dpi, mouse lungs were harvested for flow cytometry analysis, live cell sorting, and immunofluorescence staining. **(A)** Immunofluorescence analysis of CTSB protein in AM and MNC1 sorted from H37Rv-mCherry-infected mice (28 dpi). Representative images were taken by confocal microscopy. Scale bars, 10 μm. **(B)** Quantification of CTSB protein immunostaining MFI per cell for (A), >100 cells per subset were analyzed using ImageJ. **(C)** Immunofluorescence analysis of LAMP1 in AM and MNC1 sorted from H37Rv-mCherry-infected mice (28 dpi). Representative images were taken by confocal microscopy. Scale bars, 10 μm. **(D)** Quantification of LAMP1 MFI per cell for (C), >100 cells per subset were analyzed using ImageJ. **(E)** Representative histograms and MFI of intracellular LAMP1 analyzed by flow cytometry for AM and MNC1 from mice infected with Mtb H37Rv-ZsGreen (28 dpi). **(F-G)** LAMP1, CTSB and SiglecF staining of lung sections prepared from mice infected with H37Rv-mCherry (28 dpi). Scale bars, 20 μm. Green arrow indicating SiglecF^+^ AM. ****p<0.0001 by unpaired Student’s t test. Results are presented as mean ± SD, representative of 2-3 independent experiments.

We then quantitated the abundance of the lysosomal (and late endosomal) membrane protein, LAMP1 by immunofluorescence microscopy on cells sorted from lungs of infected mice. In line with the RNA-seq data, this revealed abundant LAMP1 punctate fluorescence throughout the cytoplasm of AM (Fig. 5C-5D). In contrast, LAMP1 staining of MNC1 cells was less intense, although the distribution, size, and shape of the LAMP1^+^ puncta resembled those in AM (Fig. 5C-5D). We then quantitated intracellular LAMP1 by flow cytometry and found that the MFI of intracellular LAMP1 was approximately 6-fold higher in AM than in MNC1 cells, consistent with fewer LAMP1^+^ lysosomes or lower LAMP1 content per organelle in MNC1 (Fig. 5E). These results indicate that MNC1 are deficient in classically defined lysosomes.

By immunofluorescence, the fluorescence intensity of LAMP1 and CTSB in both infected and uninfected cells within lung granulomas was lower compared with AM (Fig. 5F-5G). In addition, MNC1 retained this phenotype at 56 dpi, as seen by lower intensity staining of both LAMP1 and ATP6V1B2 compared to AM (Fig. S7). Together, these results indicate that monocyte-derived cells, especially MNC1, in Mtb-infected lungs are defective in lysosome functions as indicated by deficiencies of lysosome abundance, lysosomal protein content, and lysosomal cathepsin activities compared with AM.

### Differential expression and activation of the lysosomal regulator, TFEB, in AM and MNC1

Lysosome biogenesis and expression of genes whose products are involved in lysosome structure, acidification, and functions, are regulated by the transcription factor EB (TFEB) (*45, 46*) through recognition of CLEAR (Coordinated Lysosomal Expression and Regulation) elements (*47*).

RNA-seq of live sorted lung cell populations revealed approximately 2-fold higher TFEB expression in AM compared to MNC1, while the TFEB mRNA level was not significantly different between infected and bystander cells (Fig. 6A). TFEB localization and activity is regulated by phosphorylation, wherein phosphorylated TFEB is retained in the cytoplasm by binding to the cytoplasmic chaperone, 14-3-3, and dephosphorylated TFEB translocates to the nucleus, where it activates transcription of lysosome biogenesis genes (*48*). Immunofluorescence staining and microscopy on sorted AM and MNC1 revealed heterogeneity in the distribution of TFEB between cytoplasm and nucleus in both cell types (Fig. 6B). Despite the heterogeneity, TFEB MFI was ∼ 2 fold higher in AM compared with MNC1 (Fig. 6C). Moreover, TFEB localized to the nucleus in most, if not all, AM, while TFEB was nearly exclusively localized to the cytoplasm in MNC1 cells, a finding substantiated by quantitative image analysis of nuclear TFEB (Fig. 6D). Furthermore, the average level of TFEB of cells within lung granuloma was lower than that of AM (Fig. 6E). These results demonstrate a lower level of TFEB activity in MNC1 cells compared with AM, which is reflected downstream by expression of TFEB-regulated lysosome genes.

**Fig 6.**
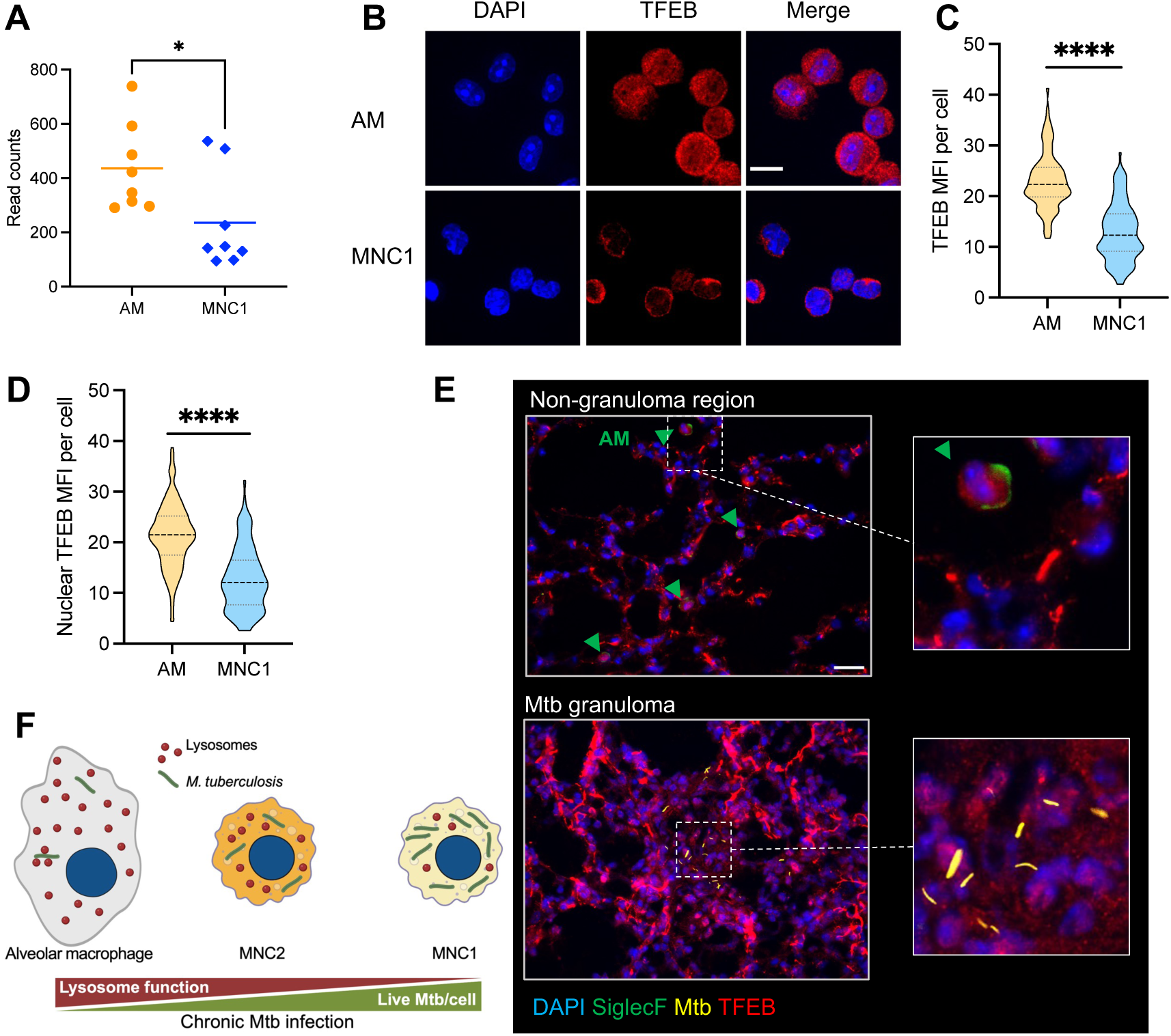
TFEB in MNC1 is predominantly extranuclear. C57BL/6 mice were infected with by aerosol with ∼100 Mtb H37Rv-mCherry. At 28 dpi, mouse lungs were harvested for live cell sorting for RNA-seq analysis, and immunofluorescence staining. **(A)** RNA-seq data evidence that MNC1 express lower levels of *Tfeb* mRNA than do AM (28 dpi). **(B)** AM have more nuclear TFEB than MNC1. Cells were isolated and sorted from lungs of mice infected with H37Rv-mCherry (28 dpi). Anti-TFEB antibody was used for detecting TFEB in sorted cells of each subset. Representative images were taken by confocal microscopy. Scale bars, 10 μm. **(C-D)** Quantification of total TFEB MFI (C) and nuclear TFEB MFI (D) per cell in (B) from >100 cells of each subset using ImageJ. **(E)** TFEB and SiglecF staining of lung sections from mice infected with Mtb H37Rv-mCherry (28 dpi). Scale bar, 20 μm. Green arrow heads indicate SiglecF^+^ AM. **(F)** A model of that Mtb restriction or survival depending on the functional lysosome abundance in distinct lung mononuclear cell subsets during chronic infection. ****p<0.0001 by unpaired Student’s t test. Data: mean ± SD, representative of 2-3 independent experiments.

### Mtb ESX-1 is required for MNC1 recruitment but does not determine MNC1 lysosome deficiency

Mtb resides in phagosomes that do not mature efficiently to phagolysosomes, and this property is dependent on the Mtb ESX-1 Type VII secretion system (*2*). Therefore, we considered the possibility that Mtb ESX-1 alters lysosome biogenesis in lung monocyte-derived cells to facilitate its persistence. To test this, we infected mice with each of three ZsGreen expressing strains: Mtb H37Rv, Mtb H37Rv:ΔRD1, and the vaccine strain *M. bovis* BCG. The latter two lack the RD1 locus encoding a key part of ESX-1. We first compared the protein levels of LAMP1 and ATP6V1B2 for subsets by measuring MFI using analytical flow cytometry. This revealed that there was no difference of LAMP1 or ATP6V1B2 MFI in MNC1 and neutrophils from naïve mice or mice infected with Mtb H37Rv, Mtb H37Rv:ΔRD1, or BCG (Fig. S8A-S8B). Interestingly, the LAMP1 and ATP6V1B2 levels of AM and MNC2 were significantly higher in Mtb H37Rv-infected mice compared with Mtb H37Rv:ΔRD1-infected mice, BCG-infected mice, or naïve mice. Despite the infection conditions, LAMP1 and ATP6V1B2 MFI of AM remained the highest among the lung phagocyte subsets. Consistent with the above results, MNC1 had lower protein levels of LAMP1 and ATP6V1B2 compared with AM, regardless of mouse infection or bacterial strain status. We further used the fluorogenic CTSB assay to assess the lysosome activities of lung cell subsets from mycobacteria-infected mice and naïve mice, where we observed results that were similar to analytical flow data for LAMP1 and ATP6V1B2 (Fig. S8C). These data suggest that the defective lysosome functions of MNC1 are not determined by Mtb ESX-1.

In contrast, we did find that Mtb ESX-1 promotes the recruitment of MNC1, MNC2, and neutrophils to the lungs, and Mtb spread from AM to these cell subsets (Fig. S8D-S8F), consistent with other results (*17, 31*). In line with these, mice infected with Mtb H37Rv:ΔRD1 or BCG showed a reduction of lung bacterial burdens compared to the mice infected with Mtb H37Rv (Fig. S8G). Together, these findings indicate that lysosome deficiency in MNC1 is not driven by Mtb ESX-1, while recruitment of lysosome-deficient permissive MNC1 is promoted by Mtb ESX-1. Based on our studies, we proposed a model in which virulent Mtb recruits and exploits lysosome-deficient lung mononuclear cells for persistence during chronic infection (Fig. 6F).

### Pharmacological activation of TFEB and lysosomal function enhances control of Mtb

Since Mtb-permissive MNC1 exhibit evidence of defective lysosomes, we investigated whether pharmacological activation of TFEB could improve restriction of Mtb replication in macrophages. We tested multiple small molecules as potential activators of TFEB in murine bone marrow-derived macrophages (BMDM) by detecting nuclear-localized TFEB (Fig. S9A-S9C). As expected, torin1, a potent inhibitor of mammalian target of rapamycin (mTOR) that regulates TFEB (*49*), induced TFEB nuclear translocation in BMDM (Fig. 7A). Consistent with reports that the c-Abl tyrosine kinase inhibitor imatinib induces lysosomal acidification to inhibit Mtb growth in human monocyte-derived macrophages (*50*) and in Mtb-infected mice (*51, 52*), we found that imatinib and an alternative c-Abl kinase inhibitor nilotinib activated TFEB nuclear translocation (Fig. 7A) to an extent comparable to that of torin1 (Fig. 7B). These results imply that c-Abl inhibitors activate TFEB and that this action accounts for certain of the earlier reports of the activity of imatinib.

**Fig 7.**
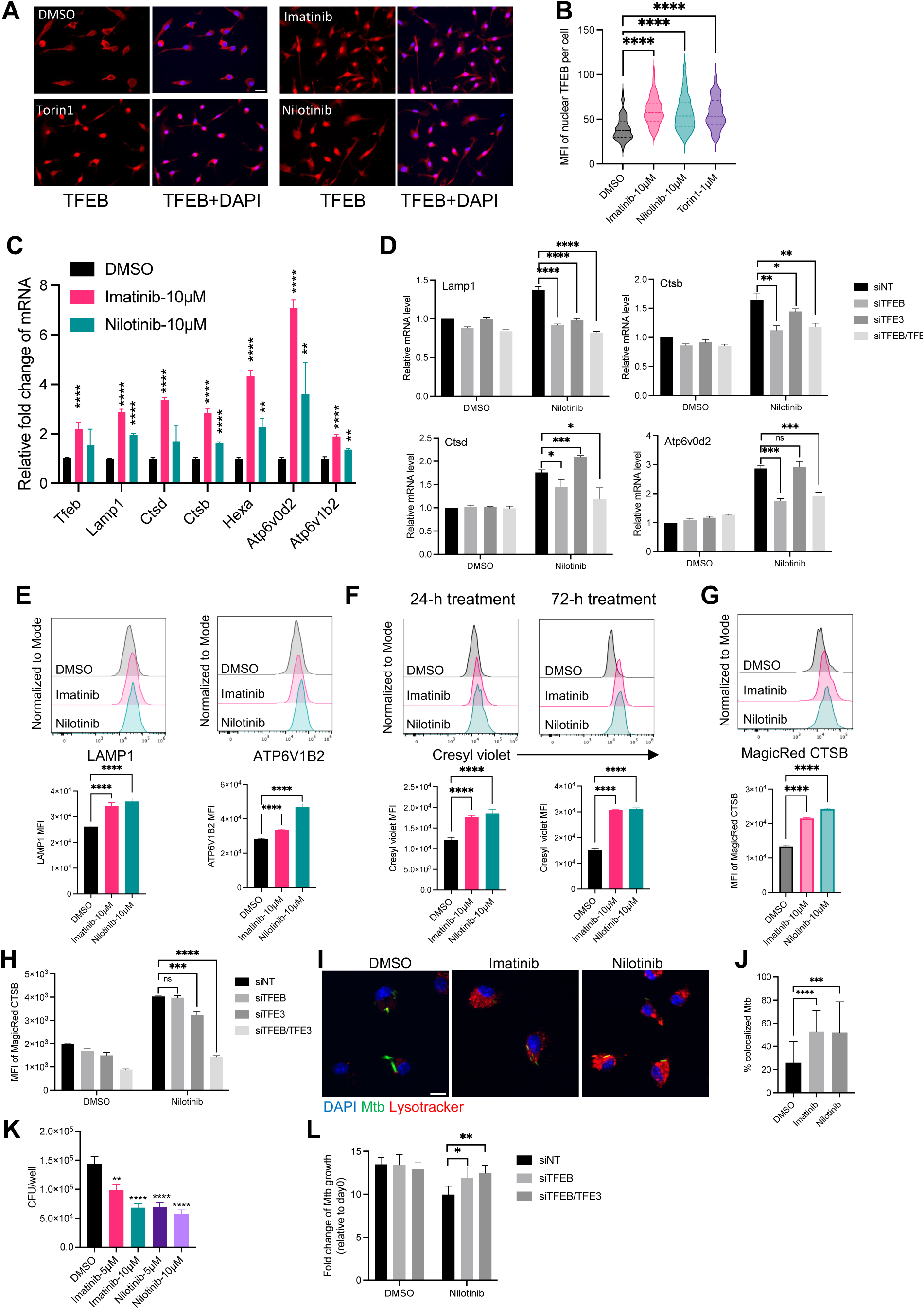
c-Abl inhibitors activate TFEB, enhance lysosome biogenesis, and improve control of intracellular Mtb in cultured primary macrophages. **(A)** Torin1 (mTOR inhibitor) and two c-Abl kinase inhibitors (imatinib and nilotinib) increase nuclear translocation of TFEB. BMDM were treated with DMSO or the indicated small molecules for 4 h. Then, cells were fixed and stained with an anti-TFEB antibody. Scale bar, 20 μm. **(B)** Quantification of nuclear TFEB MFI per cell from >70 cells for each condition in (A) using ImageJ. **(C)** Imatinib and nilotinib increase lysosome gene expression. qPCR results of lysosome genes for BMDM treated with DMSO or the indicated small molecules for 24 h. **(D)** BMDM were transfected with the designated siRNA (20 nM) for 48 h, then treated with nilotinib (5 μM) for 18 h. The mRNA levels of lysosome genes were detected by qPCR. siNT: non-targeting siRNA control. **(E)** Histograms and MFI of LAMP1 and ATP6V1B2 for uninfected BMDM treated with small molecules for 24 h. **(F)** Histograms and MFI of cresyl violet for uninfected BMDM treated with small molecules for 24 h or 72 h. **(G)** Imatinib and nilotinib induce cathepsin B enzymatic activity. Histograms and MFI of fluorogenic cathepsin B product for H37Rv-infected BMDM treated with inhibitors for 72 h. **(H)** MFI of fluorogenic cathepsin B product in siRNA transfected BMDM. **(I)** Imatinib and nilotinib enhance lysosome acidification. BMDM were infected with H37Rv-ZsGreen (MOI=2), then treated with DMSO, imatinib (10 μM), or nilotinib (5 μM). BMDM were stained with DAPI and lysotracker at 24 hpi. Scale bar, 10 μm. **(J)** Imatinib and nilotinib enhance Mtb phagolysosome maturation in cultured primary macrophages. Quantification of Mtb co-localizing with lysotracker for (G) from >20 images per condition using JACoP in ImageJ. **(K)** Imatinib and nilotinib enhance control of Mtb in primary macrophages. BMDM were infected with H37Rv at MOI=1, then treated with DMSO or indicated small molecules for 4 days. Cells were then lysed and plated for CFU assay. **(L)** siRNA-transfected BMDM were infected with Mtb H37Rv (MOI=1) and then treated with nilotinib (2 μM) for 4d. Viable intracellular bacteria were quantitated by CFU assay. Results are presented as mean ± SD, n=3-4 replicates, representative of 2-3 independent experiments. **p<0.01, ***p<0.001, ****p<0.0001 by multiple unpaired Student’s t test (C), or unpaired Student’s t test (B, D, E-I).

We then determined if c-Abl inhibitors induce expression of TFEB downstream genes using quantitative PCR (qPCR). This revealed that imatinib and nilotinib significantly increased expression of *Tfeb*, *Lamp1*, *Ctsb*, *Ctsd*, *Hexa*, *Atp6v0d2* and *Atp6v1b2* (Fig. 7C). Using siRNA knockdown, we found that induction of these lysosome genes by nilotinib mainly depends on TFEB, but not TFE3, another regulator of lysosome biogenesis in macrophages (*53*) (Fig. 7D). In line with the mRNA findings, treatment of BMDM with these TFEB activators increased LAMP1 and ATP6V1B2 proteins (Fig. 7E). We then tested if these TFEB activators induce lysosomal acidification in BMDM. We observed that imatinib or nilotinib significantly increased cresyl violet fluorescence after 24-h treatment, and further increased after 72-h treatment (Fig. 7F). In addition, CTSB enzyme activity increased in Mtb-infected BMDM treated with imatinib or nilotinib (Fig. 7G). Interestingly, the increase of CTSB activity by nilotinib depended on both TFEB and TFE3 (Fig. 7H); in agreement with this, imatinib and nilotinib also induced TFE3 nuclear translocation (Fig. S10), suggesting that TFEB and TFE3 may regulate distinct lysosome genes differently. In line with previous results in human primary macrophages (*50*), imatinib increased colocalization of Mtb with lysosomes (Fig. 7I-7J), and a similar result was obtained for nilotinib. Consistent with results of others (*50, 54*), we verified that imatinib reduced bacterial loads in Mtb-infected BMDM (Fig. 7K). We obtained similar results with nilotinib, a more specific and potent c-Abl inhibitor, indicating that the increase of lysosome activity and acidification promotes antimycobacterial activity of cultured macrophages. We found that the optimal effect of nilotinib on restricting intracellular Mtb requires TFEB and TFE3 (Fig. 7L). The antimycobacterial activity of these agents was not due to direct action on Mtb, as growth in 7H9 medium was unaffected (Fig. S9D), however we did find that they interfered with a luciferase-based assay of mycobacterial viability (not shown). Collectively, these results suggest that activation of TFEB-mediated lysosomal acidification and activity could improve the control of intracellular Mtb in macrophages.

### Nilotinib activates lysosomal functions of permissive myeloid subsets and reduces lung bacterial burden in vivo

Lysosomal activities are required for control of intracellular mycobacteria in cultured macrophages (*32–34*), and the above results and similar findings of others suggest that activation of lysosome biogenesis improves control of Mtb in macrophages (*49, 55–60*). Therefore, we asked whether c-Abl inhibitors can enhance Mtb restriction and the lysosomal functions of monocyte-derived cells in vivo (Fig. 8A). We found that nilotinib administration significantly reduced lung bacterial burden without affecting mouse body weight (Fig. 8B and Fig. S11A), in line with the previous report that imatinib treatment enhanced control of Mtb in mouse lungs (*51, 52*). Consistent with the effects in cultured macrophages, nilotinib treatment enhanced expression of multiple TFEB-activated genes in the lungs of infected mice (Fig. 8C). Flow cytometry of lung cell subsets revealed that nilotinib treatment of Mtb-infected mice significantly increased protein expression of LAMP1 and ATP6V1B2 in lung MNC1, MNC2, neutrophils and Ly6C^hi^ monocytes (Fig. 8D-8E). Nilotinib treatment also enhanced lysosome acidification in MNC1, MNC2, neutrophils and Ly6C^hi^ monocytes, but had a negligible effect on lysosome acidification in AM (Fig. 8F). We then tested the effect of nilotinib treatment on lysosome CTSB activity in these lung cell subsets and found that nilotinib significantly increased CTSB activity in MNC1, MNC2 and Ly6c^hi^ monocytes (Fig. 8G). Unexpectedly, nilotinib was associated with slightly reduced CTSB activity in AM, which still maintained higher CTSB activity than other lung cell subsets. In addition, nilotinib reduced recruitment of MNC1, MNC2, neutrophils and Ly6C^hi^ monocytes to the lungs (Fig. S11B), consistent with a secondary effect on proinflammatory responses.

**Fig 8.**
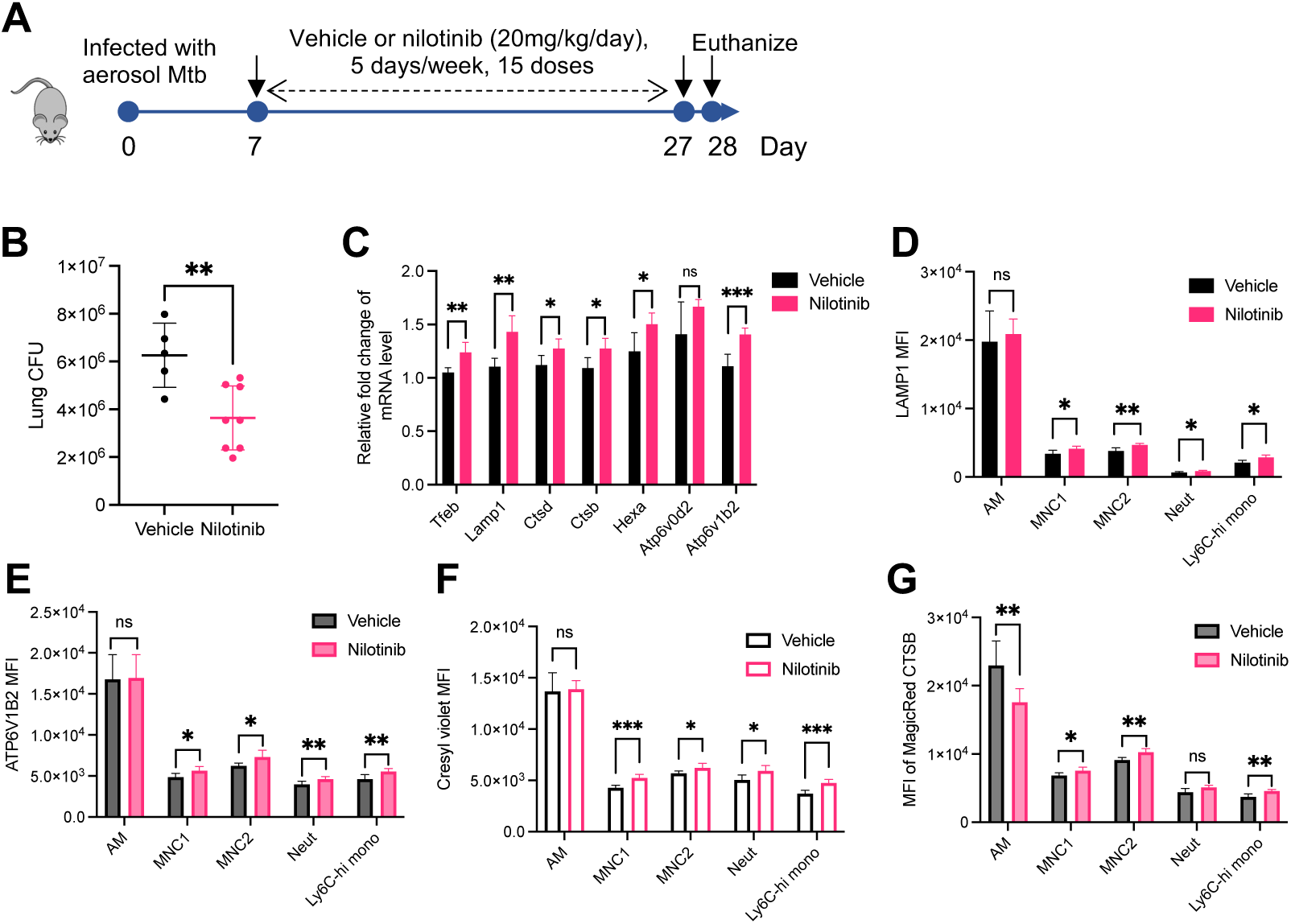
Nilotinib treatment of Mtb-infected mice activates lysosome functions of lung myeloid subsets and reduces lung bacterial burdens. **(A)** Experimental protocol for treatment of Mtb infected C57BL/6 mice. Mice infected with Mtb H37Rv-ZsGreen were treated with vehicle or nilotinib (20 mg/kg/day) intraperitoneally, 5 days/week, for a total of 15 doses beginning at 7 dpi and ending at 27 dpi. At 28 dpi, mice were euthanized, and lungs were harvested for the assays shown. **(B)** Nilotinib treatment of mice decreases overall Mtb lung bacterial burdens (28 dpi). **(C)** Nilotinib treatment of Mtb-infected mice increases expression of lysosome genes in lungs (qPCR analysis). Gapdh was used as a control. **(D)** Nilotinib treatment of Mtb-infected mice increases LAMP1 expression in recruited lung phagocyte subsets; flow cytometry quantitation of intracellular LAMP1 MFI in lung subsets from mice infected with H37Rv-ZsGreen treated with vehicle or nilotinib (28 dpi). **(E)** Nilotinib treatment of Mtb-infected mice increases lysosomal V-ATPase subunit ATP6V1B2 expression in recruited lung phagocyte subsets; flow cytometry quantitation of intracellular ATP6V1B2 MFI in lung subsets from mice infected with H37Rv-ZsGreen treated with vehicle or nilotinib (28 dpi). **(F)** Nilotinib treatment of Mtb-infected mice increases lysosomal acidification. Cresyl violet MFI quantitated by flow cytometry for lung subsets from H37Rv-ZsGreen-infected mice treated with vehicle or nilotinib (28 dpi). **(G)** Nilotinib treatment of Mtb-infected mice increases lysosomal cathepsin B activity in recruited phagocytes, but not resident (alveolar) macrophages. MFI of fluorogenic MagicRed^®^ CTSB product in lung subsets from mice treated with vehicle or nilotinib (28 dpi) quantitated by flow cytometry. Results are presented as mean ± SD, n=4-8, representative of 2 independent experiments. *p<0.05, **p<0.01, ***p<0.001, by unpaired Student’s t test (B) or multiple unpaired Student’s t test (C-G).

Activation of TFEB has been reported to enhance lysosomal proteolytic activity and increase MHCII antigen presentation (*61*). The lysosome deficiency of MNC1 suggests that they may be defective in antigen processing and presentation. By flow cytometry analysis, we observed that MNC1 express significantly lower surface MHCII compared with other mononuclear phagocytes AM and MNC2 (Fig. S11C). The MHCII level was not changed significantly by the presence of Mtb in MNC1 and MNC2, while infected AM showed increased MHCII compared to bystander AM (Fig. S11D). Interestingly, nilotinib treatment significantly enhanced the MHCII level in MNC1 and MNC2, but not AM (Fig. S11C). In addition, nilotinib treatment increased the frequency of activated (CD154^+^) CD4 T cells in the lungs, while the overall frequencies of CD4^+^ T cells, CD8^+^ T cells, and CD8^+^CD44^+^ T cells were not different than in lungs of control mice (Fig. S11E-S11G). We performed Gene Ontology (GO) analysis of differentially expressed genes in MNC1 compared with that in other subsets. This revealed multiple significant GO terms related to antigen processing and presentation (Table S1), among these, genes encoding cathepsins and MHCII are underexpressed in MNC1 compared with other subsets (Fig. S12). These results suggest that MNC1 are deficient in antigen processing and presentation, and enhancing lysosome biogenesis improves MHCII antigen processing and presentation to CD4 T cells in vivo. Collectively, these findings suggest that nilotinib rescues defective lysosomal expression and functions of permissive myeloid subsets and improves restriction of Mtb in the lungs in vivo.

## DISCUSSION

We report here that CD11c^lo^ monocyte-derived MNC1 cells are the most Mtb-permissive lung cell subset during chronic infection. Using two independent methods to quantitate live intracellular Mtb, we found that MNC1 are a major Mtb reservoir and importantly, harbor more live Mtb per infected cell than AM, neutrophils, or CD11c^hi^ MNC2 at a stage of infection when adaptive immune responses have developed, and Mtb antigen-specific effector T cells are present in the lungs. These results extend our previous finding that monocyte-derived cells are the major infected populations during chronic infection (*16*). Our results at 21 dpi are consistent with a recent report that CD11c^hi^ monocyte-derived cells (MNC2 in this study) are the major infected subset at the same timepoint (*12*), yet they differ from those results at other time points, at least in part because we specifically quantitated live intracellular bacteria. Our results emphasize that Mtb infection is highly dynamic, especially before and during the development of adaptive immune responses, and that distinct lung myeloid cell subsets can prevail as niches for Mtb at different stages of infection.

Previous studies have shown that lysosomal enzymes are required for antimycobacterial activity (*32–34*). In addition, monocyte-derived macrophages from TB resisters (TB contacts who developed neither asymptomatic LTBI nor active TB) exhibit better Mtb killing ability via phagosome acidification compared to healthy controls, latent TB controls, and active TB patients (*62*). We found that MNC1 are deficient in lysosome functions due to low lysosome content, poor lysosomal acidification and reduced enzyme activity compared to AM, and lysosome functions in these lung cell subsets correspond to their ability to restrict intracellular Mtb 28 dpi (Fig. 6F). Lysosome deficiency of MNC1 is not due to a direct effect of Mtb or the essential virulence factor ESX-1 on individual cells. Instead, Mtb induces the recruitment of permissive MNC1 and less permissive MNC2 to the lungs in an ESX-1 dependent manner. Therefore, in addition to blocking cell autonomous phagosome maturation and autophagy (*2, 63*), Mtb exploits lysosome-poor monocyte-derived cells for persistence during chronic infection. Lysosome function is essential for autophagy, which is also required for restricting Mtb (*64–68*). However, whether autophagy function differs between AM and monocyte-derived cells remains to be determined.

TFEB is a master transcription factor for lysosome biogenesis (*46*). Compared with AM, MNC1 have less nuclear TFEB (i.e., activated TFEB) and lower expression of TFEB-activated genes at both the mRNA and protein levels. This implies that TFEB activation regulates the difference in lysosome content and function between these subsets. Notably, TFEB is required for IFNγ-dependent control of Mtb in cultured macrophages (*49*). TFEB activators can boost the antimycobacterial activity of macrophages. These activators include bedaquiline (*55*), NR1D1 agonist (*69*), PPARα agonist (*56*), trehalose (*70*), and IFNγ (*58*). We found that the c-Abl tyrosine kinase inhibitors imatinib and nilotinib activate TFEB and induce expression of downstream genes to improve lysosome function in macrophages. Like imatinib, nilotinib enhanced control of Mtb in vitro and in vivo, suggesting enhancement of lysosome function can partially overcome the reported inhibitory effects of Mtb on phagosome maturation. In vivo, nilotinib improved lysosome functions in MNC1, MNC2, and Ly6C^hi^ monocytes, suggesting the potential for enhancing lysosome functions in these Mtb-permissive monocyte-derived cells as a host-directed therapy for TB. TFEB-mediated lysosome biogenesis is also important for autophagy (*46*). Like other TFEB activators, nilotinib has been reported to activate autophagy in macrophages (*71*). Therefore, nilotinib and other TFEB activators likely improve Mtb restriction in part via enhancing phagosome maturation and autophagy.

We found evidence that MNC1 are defective in antigen processing and presentation. This may partly be due to lysosome deficiency, as TFEB-mediated lysosomal proteolytic activity increases MHCII antigen presentation in antigen-presenting cells (*61*). These results are consistent with our prior observation that MNC1 (then termed recruited macrophages) are inferior to AM and MNC2 (then termed monocyte-derived dendritic cells) in activating Mtb antigen-specific CD4 T cells (*72*). Consistent with nilotinib enhancement of MHCII expression and antigen presentation through increased lysosome activities, we found increased frequencies of activated CD4 T cells in lungs of mice treated with nilotinib. Therefore, increasing lysosome functions in lung mononuclear cells may enhance control of Mtb by multiple mechanisms in vivo.

In this work, the effect of nilotinib on lung bacterial burdens was modest. Possibly because Mtb blocks phagosome maturation even in the face of increased lysosome biogenesis; however, it is also possible that the limited effect is due to suboptimal pharmacokinetics or off-target effects that counteract the positive effects.

Our study supports that the roles of distinct lung mononuclear cell subsets evolve during Mtb infection. AM serve as an early replication niche for Mtb (*9, 14*), and are anti-inflammatory and permissive compared to IM in early innate immune responses to Mtb (*11, 14*), due to induction of an NRF2-dependent cell-protective antioxidant response (*14*). However, AM become pro-inflammatory and upregulate IFNγ-responsive genes after the development of adaptive immune responses (*12, 27*). Our studies also revealed that IFNγ signaling is upregulated in AM compared to MNC1, as indicated by higher mRNA levels of Ifngr2, H-2 and Ciita (Data file S1), and a previous study revealed that IFNγ signaling reprograms AM to better control Mtb growth (*73*). In line with this, the surface MHCII level of AM from Mtb-infected mice (28 dpi) is ∼13 fold higher than that of AM from naïve mice (data not shown). Since IFNγ can activate TFEB-mediated lysosome biogenesis (*58*), we hypothesize that improved lysosome function via IFNγ signaling contributes to better Mtb killing activity in AM during chronic infection. Thus, during chronic infection, AM become Mtb restrictive partially via IFNγ signaling while MNC1 remain permissive despite exposure to IFNγ.

Although certain lysosome genes are highly expressed in neutrophils compared to MNC1, we did not find neutrophils have higher lysosome functions than MNC1, suggesting that different mechanisms regulate lysosome activities in neutrophils. Unlike mononuclear phagocytes, neutrophils do not have a classic endosomal pathway or classical lysosomes. Instead, they possess lysosome-like granules containing bactericidal factors that can rapidly fuse with phagosomes (*74*). In addition, neutrophil phagosomes have neutral pH, and LAMP proteins are absent from their granules (*74, 75*). Thus, neutrophils use mechanisms distinct from those of mononuclear phagocytes to kill ingested pathogens, and these mechanisms (including NADPH oxidase activity) are not correlated with lysosome function.

A limitation of this study is the possibility that our cell subsets are heterogeneous, partially due to the lack of discriminating surface markers and limited fluorescence parameters of our BSL3-contained cell sorter. Although single-cell RNA sequencing has led to identifying other monocyte-derived cell subsets during Mtb infection (*27*), discriminating surface markers for better separation of these subsets still need improvement. Nevertheless, the resolution of our flow sorting strategy was sufficient to identify four transcriptionally distinct phagocyte subsets, including MNC1 that harbor the most live bacteria compared to the others. Further translational work will be required to ascertain lysosome function in lung mononuclear subsets in Mtb-infected humans; such studies face the challenges of obtaining lung parenchymal cells (and not only AM by bronchoalveolar lavage) from people with active tuberculosis.

In summary, our work revealed that Mtb recruits monocyte-derived MNC1 that are intrinsically deficient in lysosome biogenesis and that enable Mtb persistence during chronic infection. The c-Abl inhibitor nilotinib activates TFEB and improves lysosome functions of monocyte-derived cells including MNC1, leading to enhanced control of Mtb in vitro and in vivo. Fig. 6F shows a graphic depiction of our working model based on results of our present studies. Inhibiting c-Abl to enhance lysosome biogenesis represents a promising strategy for developing host-directed therapeutics to reprogram permissive cells to better control Mtb. In addition, understanding the mechanisms by which restrictive subsets combat Mtb infection will also provide valuable insight into the mechanisms that limit protective immunity to Mtb.

## MATERIALS AND METHODS

### Study design

This study aimed to characterize the lung myeloid subsets that serve as a permissive niche for Mtb during chronic infection, and to identify mechanisms that enable Mtb to persist. All experiments used randomly assigned mice. The number of biological replicates and the number of repetitions for each experiment are indicated in figure legends.

### Chemicals

Chemicals used in this study were from the following sources: nilotinib (Selleckchem, S1033), imatinib (Selleckchem, S2475), torin1 (Selleckchem, S2827), Gefitinib (MedChemExpress, HY-50895), GSK621 (Sigma, SML2003), EX229 (MedChemExpress, HY-112769), BC1618 (MedChemExpress, HY-134656), bafilomycin A1 (Selleckchem, S1413), MK2206 (Selleckchem, S1078), SRT1720 (Apexbio, A4180-10), FR180204 (Selleckchem, S7524), PLX-4720 (Selleckchem, S1152). PEG300 (Selleckchem, S6704), DMSO (Sigma, D2650).

### Mice

All animal experiments were approved by the Institutional Animal Care and Use Committee of University of California, San Francisco or NYU School of Medicine. 8-week-old C57BL/6 mice were obtained from Jackson Laboratory.

### Bacterial strains and growth

Mtb H37Rv was transformed with plasmid (pMV261::EGFP, pMV261::ZsGreen or pMSP12-mCherry) to constitutively express the respective fluorescent protein. H37Rv-live/dead was generated by transforming Mtb with the Live/Dead plasmid that drives constitutive expression of mCherry and tetracycline-inducible GFP (*30*). BCG Pasteur was transformed with pMV261::ZsGreen. Bacteria were grown in Middlebrook 7H9 medium (BD) supplemented with 10% (v/v) ADC (albumin, dextrose, catalase), 0.05 % Tween 80, 0.2 % glycerol and 50 μg/ml kanamycin (for recombinant stains carrying pMV261 or pMSP12 plasmid).

### Generation and infection of BMDM

To generate BMDM (*72*), bone marrow cells were cultured in BMDM medium [DMEM (Gibco, 11965092), 10% heat-inactivated FBS (HI-FBS), and 20 ng/ml recombinant murine M-CSF (PeproTech)] for 6 days. Before infection, cells were washed twice with PBS and harvested with PBS/2mM EDTA, resuspended in BMDM medium. 0.510×^5^ BMDM were seeded in 96-well plates. To prepare single-cell Mtb suspensions, 5mL mid-log Mtb cultures were centrifuged at 3500rpm for 5min, washed in 5 mL DMEM/10% HI-FBS, and resuspended in 5mL BMDM infection media. The Mtb suspension was centrifuged at 1000rpm for 3min. 3mL of supernatant was transferred to a new tube and OD_600nm_ was measured. Inoculum was prepared by diluting the supernatant with BMDM media. Media were aspirated and 100 μL of inoculum was added to the well. After incubation for 3h (37°C, 5% CO_2_), cells were washed three times with DMEM/1% HI-FBS and incubated in 200 μL of BMDM media containing small molecules of interest. After 4 days, medium was removed, and cells were lysed with 100 μL 0.1% Triton X-100 in sterile H_2_O for 5-10 min. Lysates were serially diluted in PBS/0.05% Tween 80, 50 μL of sample was plated on 7H11 agar plates. CFU were counted 3 weeks later.

### Aerosol infection

Mice were infected with Mtb or BCG via aerosol using an inhalation exposure unit from Glas-Col as previously described (*13, 16, 22, 25, 26, 72, 76-86*). For controls of fluorescent protein expressing Mtb, mice were infected with H37Rv by the same procedure on the same day. Target dose was ∼100 CFU/mouse for wild type Mtb, ∼600 CFU/mouse for Mtb ΔRD1 strain, respectively. For BCG aerosol infection, we used a higher dose ∼210×^4^ CFU/mouse, as lower doses lead to bacterial clearance. Infection dose was determined by plating lung homogenates 24 hpi on 7H11 agar plates, and counting CFU after 3 weeks incubation at 37 °C. For induction of GFP expression in mice infected with live/dead-H37Rv, 1 mg/mL of doxycycline was given to mice via drinking water containing 5 % sucrose for 6 days before day 28 harvest.

### Lung homogenate preparation

Lung homogenates were prepared as previously described with modifications (*13*). Lungs were perfused with 10 mL of PBS/2 mM EDTA via right ventricle immediately after euthanasia. Lungs were chopped into small pieces, minced with a gentleMACS dissociator (Miltenyi, lung program1), and digested in 4 mL of RPMI-1640/5% HI-FBS containing 1 mg/mL collagenase D (Sigma-Aldrich) and 30 μg/mL DNase I (Sigma) for 30 min at 37 °C. Digested lung tissues were further minced with the gentleMACS dissociator (lung program2) then passed through a 70-μm cell strainer. Red blood cells were lysed with 3mL ACK lysis buffer (Gibco) for 3 min and washed twice with RPMI-1640/5% HI-FBS. Lung cells were further processed as needed.

### Flow cytometry and cell sorting

Lung cells prepared as described above were counted and washed with cold PBS twice before staining with 1:200 Zombie Aqua Fixable Viability Dye (BioLegend, 423101) for 15 min at 4 °C, and then blocking with 1:100 CD16/CD32 (BD, 553142) for 10 minutes. Leaving the blocking antibody in, the cells were then stained with the antibodies diluted in Brilliant Stain Buffer (BD, 566349). Antibodies for surface makers of myeloid cells were: Ly6G-BV421 (BioLegend, 127628), CD11c-BV605 (BioLegend, 117334), CD11b-BV711 (BioLegend, 101241), CD90.2-PE-Cy5 (BioLegend, 105314), CD19-PE-Cy5 (BioLegend, 115510), NK1.1-PE-Cy5 (BioLegend, 108716), Ly6C-PE-Cy7 (BioLegend, 128018), I-A/I-E (MHCII)-Alexa Fluor 647 (BioLegend, 107606), SiglecF-APC-Cy7 (BD Biosciences, 565527), I-A/I-E (MHCII)-PE (BioLegend, 107608), MerTK-PE (eBioscience, 12-5751-80), CD64-PE/Dazzle594 (BioLegend, 139320) and F4/80-StarBright Violet 570 (BIO-RAD, MCA497SBV570). Antibodies for T cell panel were: CD4-BV605 (BioLegend, 100548), CD8a-BV650 (BioLegend, 100742), CD69-PerCP/Cy5.5 (BioLegend, 104522), CD44-PE/Cy7 (BioLegend, 103030), CD154-APC (BioLegend, 106510), CD3-Alexa Fluor 700 (BioLegend, 100216). Antibodies for intracellular markers were: CD107a (LAMP1)-Alexa Fluor 594 (BioLegend, 121622), ATP6V1B2-Alexa Fluor 647 (Santa Cruz, sc-166045). Surface staining was performed for 30 min at 4°C, and samples were further processed as below. For flow cytometry analysis, stained cells were washed twice with PBS and fixed overnight in 1% paraformaldehyde (PFA). Samples were analyzed using a BD LSRII or Sony MA900.

For live cell sorting, samples were washed twice with FACS buffer (PBS+2mM EDTA + 2% heat-inactivated FBS), resuspended in FACS buffer and passed through a 50 μm strainer (concentration 10-2010×^6^ cells/mL). Cell subsets were sorted using a Synergy cell sorter or Sony MA900 cell sorter through 100 μm nozzle. For intracellular staining, cells were fixed and permeabilized using BD Fixation/Permeabilization Kit for 20 min at 4°C after surface staining, then washed and incubated with antibodies diluted in 1x BD Perm Wash/0.5% FBS for 30 min at room temperature. Samples were washed and acquired using a BD LSRII or Sony MA900.

### Bacterial quantitation in sorted cells

For CFU of infected cells, 1000 infected (mCherry^+^ or EGFP^+^) cells from each subset were sorted into RPMI 1640/5% HI-FBS, spun down and resuspended in 50 uL of 0.5% Tween 80/PBS. Serial dilutions were made using 0.05% Tween 80/PBS and plated on 24-well 7H11 agar plates. CFU were counted after 3 weeks. For quantitation of live and dead Mtb per cells, mCherry^+^ cells were sorted into RPMI 1640/5% HI-FBS, spun down onto Shandon Cytoslide and fixed in 1 % PFA overnight. Live (mCherry^+^GFP^+^) or dead (mCherry^+^GFP^-^) Mtb per infected cell (≥ 300 cells/condition) were quantitated.

### RNA sequencing and data analysis

Ten thousand cells of each infected or bystander subset were sorted into RNAlater and stored at -20°C until use. Total RNA was extracted using RNeasy Plus Mini Kit (QIAGEN), and libraries were prepared by polyA selection using oligo-dT beads (Life Technologies) which were sequenced on the Illumina HiSeq 2500 following standard protocols to achieve 50 nucleotide, paired end reads.

RNA sequence processing and alignment: FastQC2 (v0.11.9) was used to generate quality-control reports of individual FASTQ files. Read were then aligned to the Mus musculus (house mouse) genome assembly GRCm38 (mm10) with the splice-aware STAR aligner using ‘GeneCounts’ quant mode with the Ensembl gene transfer format file (indexes were built using ‘—sjdbOverhang 50’ and the gtf file). Biotypes were investigated using PICARD tools (v2.22.3).

Differential gene expression and pathway analysis: Differential gene expression analyses were done in R using the DESeq2 package (v1.26.0) which uses a negative binomial generalized linear model. Blinded variance stabilizing transformation was applied and t-sne plots were then generated with Rtsne (v0.15) using PCA and a perplexity of three. Differentially expressed genes were identified using a Benjamini-Hochberg corrected alpha of 0.05 and an absolute effect size of one. Kyoto Encyclopedia of Genes and Genomes (KEGG; v4.0) pathway analyses were done using the R package clusterProfiler (v3.14.3). Analyses of custom pathways considering the gene background and not were done using camera and roast (using 99990 rotations) from the limma package (v3.42.2) respectively.

### Cathepsin enzymatic activity assays

Magic Red^®^ Cathepsin (B, L or K) assay kit (Bio-Rad) was used to determine the activity of cathepsin in cells. For immunofluorescence, subsets were sorted into RPMI1640/10% HI-FBS. 50,000 cells were plated in 8-well Nunc Lab-Tek chamber slides (Thermo Scientific, 177445), incubated in cell incubator for 1h. 200μL of media containing CTSB substrate (1:250 dilution) or CTSB+CTSL+CTSK substrates (1:750 dilution for each substrate) were added and incubated for 1h at 37°C. Control samples were treated with the substrates in the presence of bafilomycin A1 (100 nM).Cells were washed 2 times and fixed in 1% PFA overnight at 4 °C. Images were taken using an in-house Leica fluorescent microscope or a CSU-22 Spinning Disk Confocal at UCSF Center for Advanced Microscopy.

For flow cytometry analysis, 2-310×^6^ lung cells or 0.510×^6^ BMDM in 24-well plates were incubated with CTSB substrate for 30 min at 37 °C. Lung cells were washed twice in PBS and were processed according to the above protocol for flow cytometry. BMDM were washed and harvest with Cellstripper buffer. Data were acquired on a Sony MA900.

### Staining of acidic lysosomes

Cresyl violet (Sigma) or LysoTracker DND-99 (Thermo Fisher) was used to label acidic lysosomes. For fluorescence analysis of sorted subsets or BMDM, cells were incubated with 5 μM cresyl violet or 200 nM LysoTracker DND-99 for 1 h at 37 °C. Cells were washed 2 times and fixed in 1% PFA overnight at 4 °C. Images were taken using an in-house Leica fluorescent microscope or a CSU-22 Spinning Disk Confocal. For flow cytometry, cells were incubated with 2 μM cresyl violet for 30 min at 37 °C and processed as described in the cathepsin activity assays.

### Immunofluorescence

Lung sections were prepared as described previously (*13*). Primary antibodies used in this study were: anti-TFEB (Bethyl Laboratories, A303-673A), anti-LAMP1 (Invitrogen, 14-1071-82), anti-CTSB (Proteintech, 12216-1-AP), anti-ATP6V0D2 (Sigma, SAB2103221), anti-F4/80 (eBioscience, 14-4801-81), anti-CD68 (Bio-Rad, MCA1957T). Anti-CTSB (CA10) was also used for detecting CTSB in sorted subsets. For rabbit primary antibodies, tissue sections were incubated with 100 μL of primary antibody (1:200) in PBS + 5% goat serum for 1-1.5 h at room temperature, followed by staining with 1:500 goat anti-rabbit IgG (H+L) secondary antibody tagged with Alexa Fluor 647 (Invitrogen, A-21244) + 1:50 SiglecF-Alexa Fluor 488 (BD, 567005) + 1:100 mouse Fc block (BD) for 1h at room temperature. Slides were washed 3 times with PBS and mounted with ProLong Diamond Antifade Mountant with DAPI. For rat primary antibodies, tissue sections were stained with 100 μL of primary antibody (1:200) for 1-1.5 h, followed by incubation with goat anti-rat IgG (H+L) secondary antibody tagged with Alexa Fluor 488 (Invitrogen, A-21470). Tissue sections were washed and blocked with 1:100 mouse Fc block in 5% goat serum, then stained with 1:50 CD11b-Alexa Fluor 647 (BioLegend, 101218) for 1 h. Slides were washed and mounted with ProLong Diamond Antifade Mountant with DAPI.

For immunofluorescence staining of BMDM or sorted cells, fixed cells were permeabilized with 0.1% Triton X-100 for 10 min and blocked with 5% goat serum for 30min. Cells were then incubated with 1:500 primary antibody in 5% goat serum, washed and stained with 1:1000 secondary antibody tagged with Alexa Fluor 647. Samples were washed and mounted ProLong Diamond Antifade Mountant with DAPI.

### Quantitative PCR (qPCR)

110×^6^ BMDM were seeded in 6-well plates overnight, then treated with small molecules according to the specific experiment. After 24 h, medium was removed, and cells were lysed with 350 μl trizol (Invitrogen). Total RNA was extracted using a Direct-zol RNA Microprep kit with DNase I (R2062). 500 ng of RNA was used for cDNA synthesis using a PrimeScript RT Reagent Kit (Takara, RR037A). 2 μL of diluted cDNA (1:50) was used as template for qPCR reaction using PowerUp SYBR Green Master Mix (Applied Biosystems, A25742). For internal control, Gapdh was used. Primers were listed in Table S2.

### siRNA transfection

siRNA transfection was performed using Lipofectamine™ RNAiMAX Transfection Reagent according to the manufacture’s instruction. BMDM were reversely transfected with ON-TARGETplus SMARTpool siRNA (20 nM) targeting mouse TFEB (Dharmacon, L-050607-02-0005) or TFE3 (Dharmacon, L-054750-00-0005), and a non-targeting control siRNA pool (Dharmacon, D-001810-10-05). After 48 h, cells were used for experiments.

### Quantification and statistical analysis

Quantification of MFI from immunofluorescence images was done using ImageJ. Flow cytometry data were analyzed using FlowJo (version 10.8.1). GraphPad Prism software (version 9) was used for graphical presentation and statistical analysis. Results were presented as means ± SD. unpaired Student’s t test was done to compare two groups. For more than two experimental groups, we used one-way analysis of variance (ANOVA) or two-way ANOVA for one or two variable analysis, respectively. P > 0.05 was considered not significant (ns); P < 0.05 was considered significant; *P < 0.05; **P < 0.01; ***P < 0.001; ****P < 0.0001.

## SUPPLEMENTARY MATERIALS

Figs. S1 to S12

Table S1 to S2

Data file S1

## Acknowledgments

We thank Dr. Jessica Jang for her contributions to initial experiments in this project, and Dr. Fei Ning for her assistance in mouse cell harvests. We also thank Dr. Samuel Behar for providing the live/dead reporter plasmid for Mtb.

## Author contributions

W.Z. and J.D.E. designed the research. J.D.E. supervised the study. W.Z., I.C., J.M.B., B.S.Z., and Z.H. performed the experiments. J.L. performed RNA-seq analysis and prepared the related figure and table. W.Z. and J.D.E. wrote the manuscript, the other authors approved/edited the manuscript.

