## Supplemental Files for "*Mycobacterium tuberculosis* resides in lysosome-poor monocyte-derived lung cells during chronic infection"

Weihao Zheng *et al.*

**The PDF file includes:**
Figs. S1 to S12
Table S1 to S2

**Other Supplementary Material for this manuscript includes the following:**
Data file S1

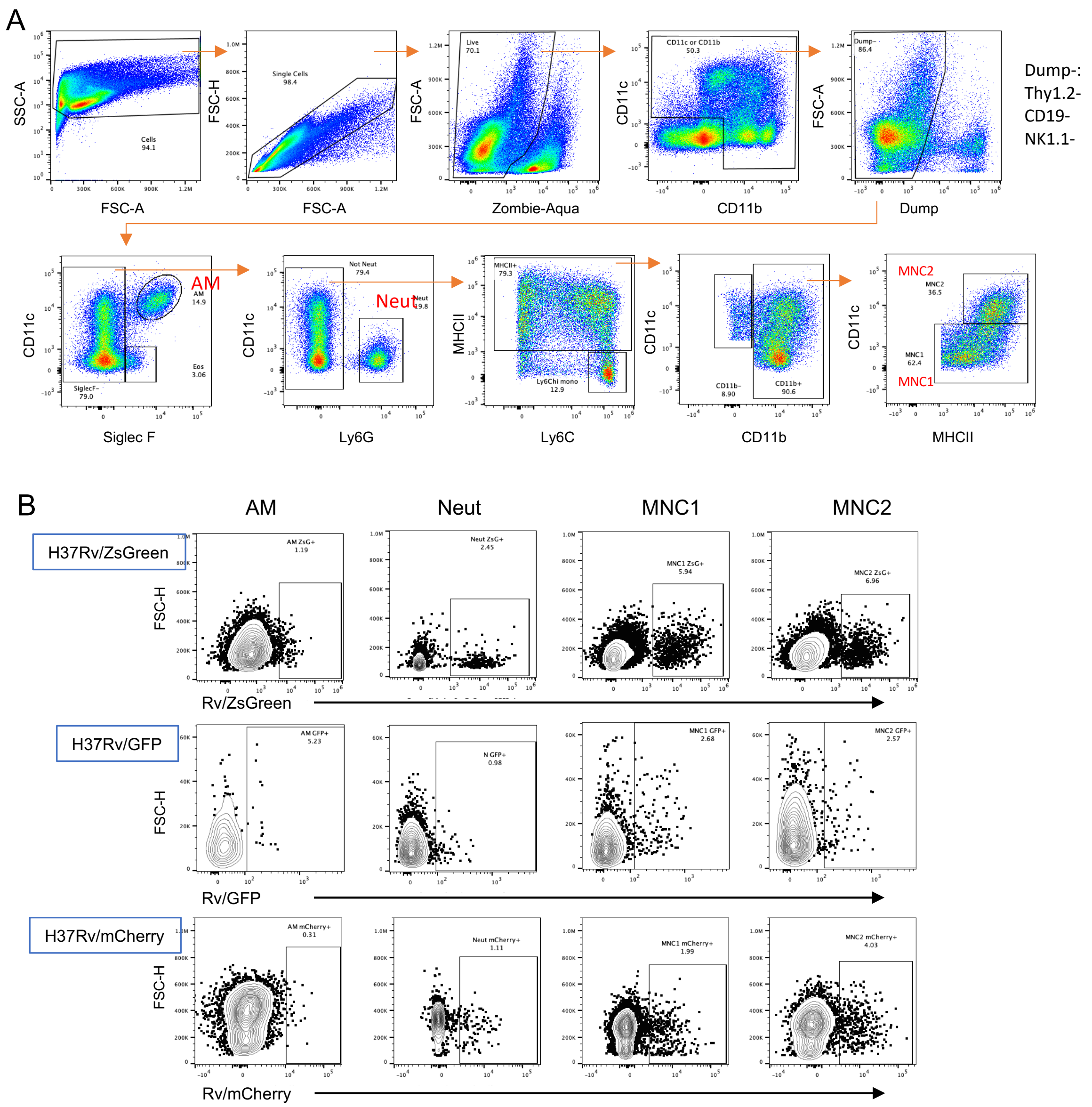


**Fig. S1. Gating strategy used to identify myeloid subsets and infected cells.**

(**A**) The designed flow panel to detect AM, neutrophil (N), MNC1, and MNC2 populations in lungs from Mtb-infected mice. After gating out B, T, and NK cells, AM were CD11b$^{lo}$CD11c$^{hi}$SiglecF$^{hi}$, MNC1 were SiglecF$^{-}$CD11b$^{+}$CD11c$^{lo}$MHCII$^{+}$, MNC2 were SiglecF$^{-}$CD11b$^{+}$CD11c$^{hi}$MHCII$^{hi}$, and neutrophils (N) were SiglecF$^{-}$Ly6G$^{hi}$CD11b$^{hi}$.

(**B**) Illustrative plots of infected lung cells in each subset from mice infected with H37Rv-ZsGreen, H37Rv-GFP, or H37Rv-mCherry (28 dpi).

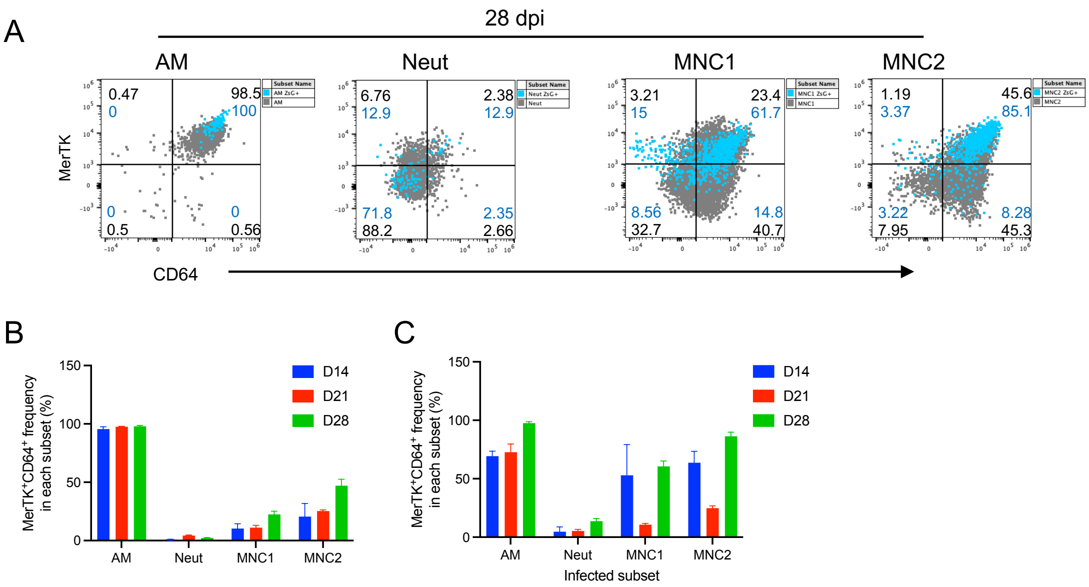


**Fig. S2. Use of MerTK and CD64 criteria excludes major fractions of recruited monocyte-derived cells and Mtb-infected cells during Mtb infection.**

(**A**) Representative plots of MerTK and CD64 expression on lung subsets (grey dots), or infected cells (blue dots) from each subset of mice infected with H37Rv-ZsGreen (28 dpi). MerTK and CD64 were added to the flow panel shown in Fig. S1A.

(**B**) MerTK$^{+}$CD64$^{+}$ frequency in each subset from mice infected with H37Rv-ZsGreen for 14-28 days of infection.

(**C**) MerTK$^{+}$CD64$^{+}$ frequency of infected cells in each infected subset from mice infected with H37Rv-ZsGreen for 14-28 days of infection.

Results are presented as mean ± SD of 4-5 mice.

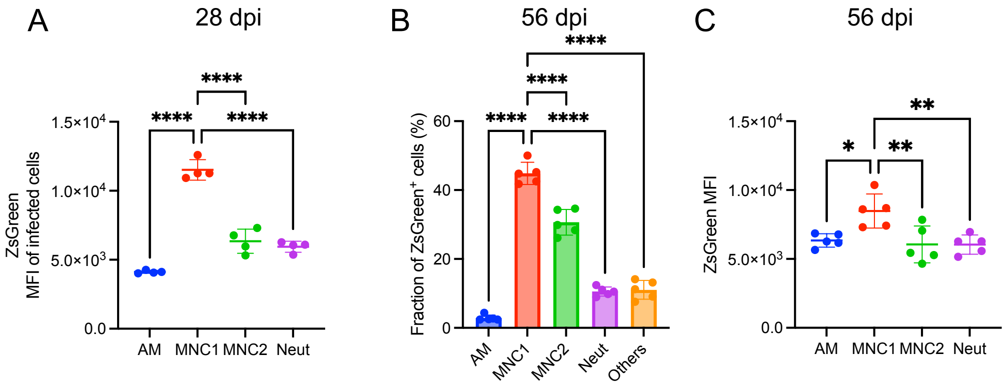


**Fig. S3. MNC1 harbor more bacteria at a later stage of chronic Mtb infection.**
C57BL/6 mice were infected with low-dose aerosolized Mtb. At 28 dpi or 56 dpi, mouse lungs were harvested for flow cytometry analysis.
(**A**) related to Fig 1A-1C. ZsGreen MFI of infected subsets from mice infected with H37Rv-ZsGreen (28 dpi).
(**B**) Flow cytometry was used to analyze the subset faction of infected cells in mouse lungs infected with H37Rv-ZsGreen (56 dpi).
(**C**) ZsGreen MFI of infected subsets from mice infected with H37Rv-ZsGreen (56 dpi).
Results are presented as mean ± SD of 4-5 mice. *p<0.05, **p<0.01, ****p<0.0001 by one-way ANOVA.

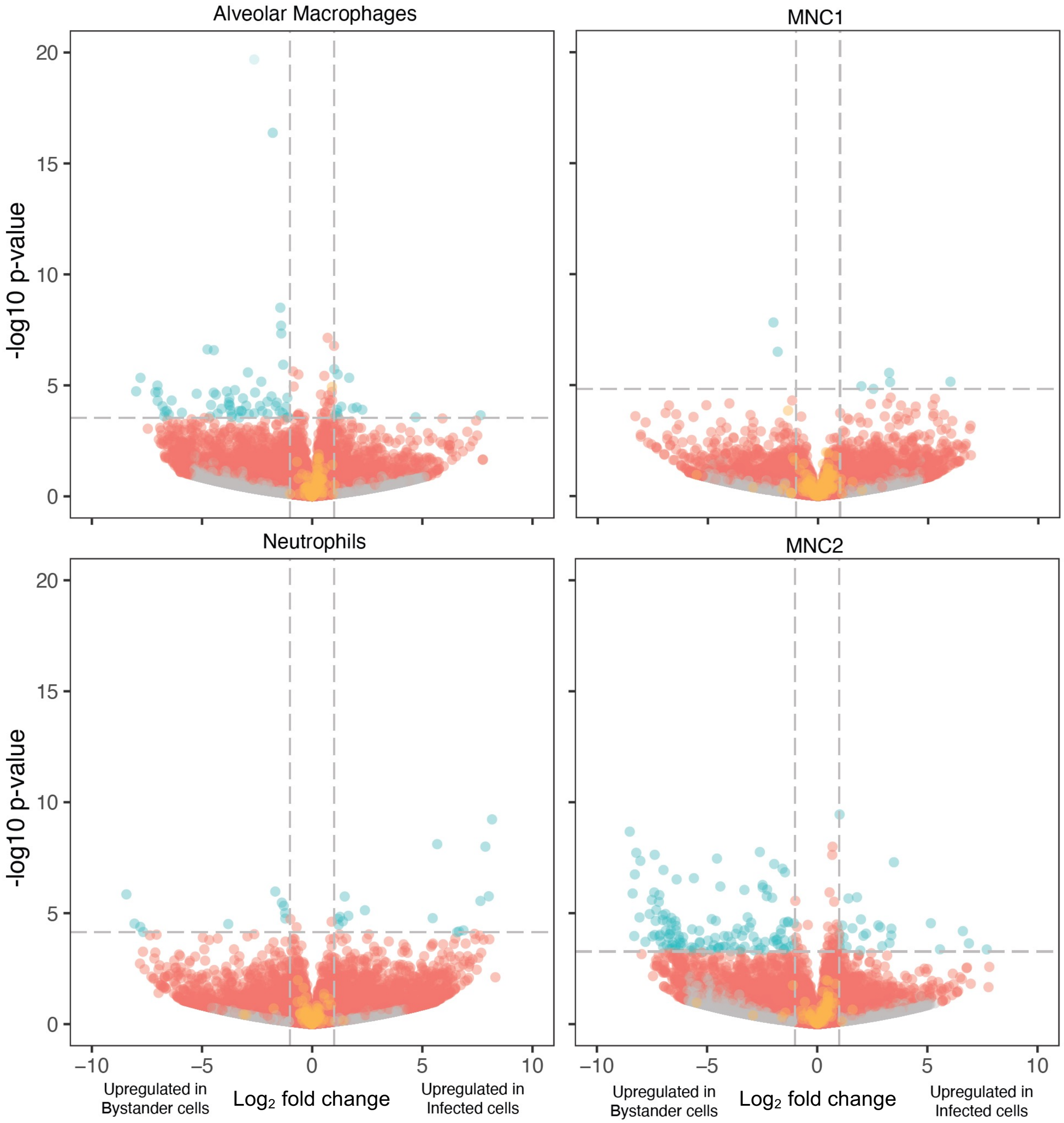


**Fig. S4. Volcano plots showing differentially expressed genes between infected cells and bystander cells for each lung cell subset.** The green dot indicates significant genes with an adjusted p-value ≤ 0.05 and a |log2 fold change| ≥ 1, the red dot indicates non-significant genes, and the grey dot indicates genes filtered out of the analysis based on the Crooks index.

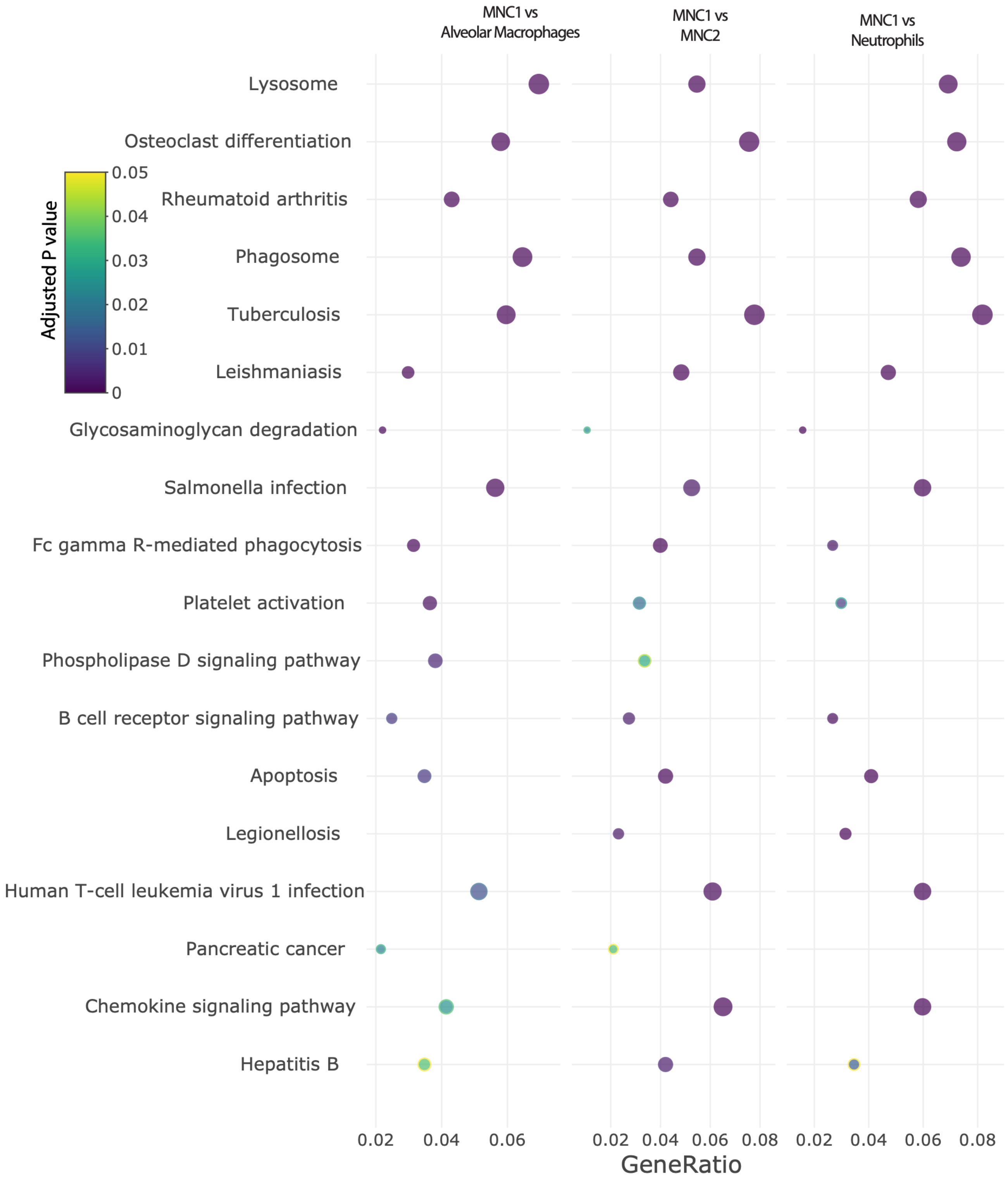


**Fig. S5. Dot plot showing 18 KEGG pathways that differ significantly with an enrichment ratio greater than 0.04 for AM, MNC2, neutrophils, and MNC1.** The color represents the adjusted p values, the graph is ordered by descending values for MNC1 vs AM, while the dot size is proportional to the gene count.

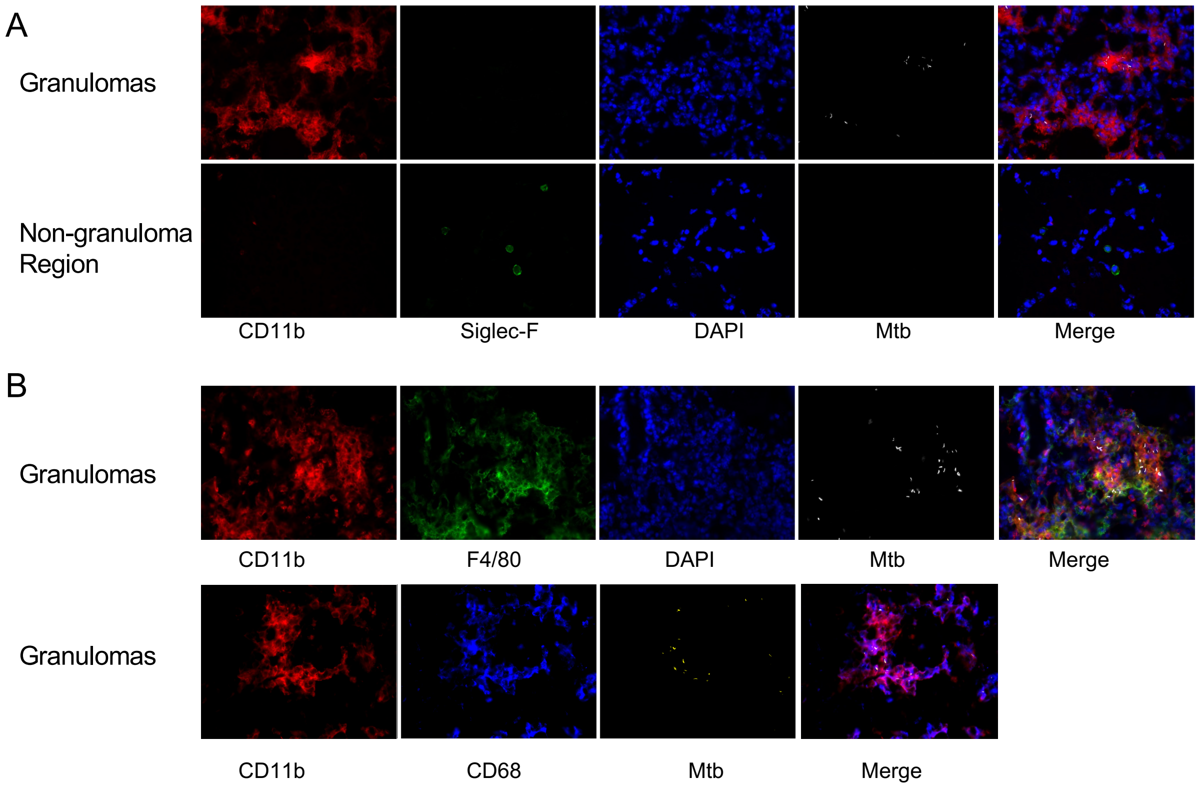


**Fig. S6. AM are excluded from granulomas, and are common in non-granuloma/uninvolved regions of the lungs.**

(**A**) Lung sections from mice infected with H37Rv-mCherry (28 dpi) were stained with CD11b-Alexa Fluor 647 and SiglecF-Alexa Fluor 488.

(**B**) F4/80 and CD68 colocalize with CD11b in lung sections prepared from mice infected with H37Rv/mCherry (28 dpi).

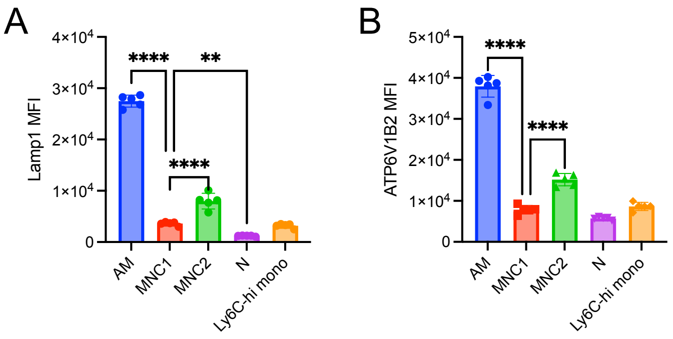


**Fig. S7. MNC1 are deficient in lysosomal proteins at a later stage of chronic Mtb infection (56 dpi).**

C57BL/6 mice were infected with low-dose aerosolized Mtb H37Rv-ZsGreen. Mouse lungs were harvested for flow cytometry analysis at 56 dpi.

(**A**) LAMP1 MFI of lung subsets from H37Rv-ZsGreen-infected mice (56 dpi).

(**B**) ATP6V1B2 MFI of lung subsets from H37Rv-ZsGreen-infected mice (56 dpi).
Results are presented as mean ± SD of 4-5 mice. **p<0.01, ****p<0.0001 by one-way ANOVA.

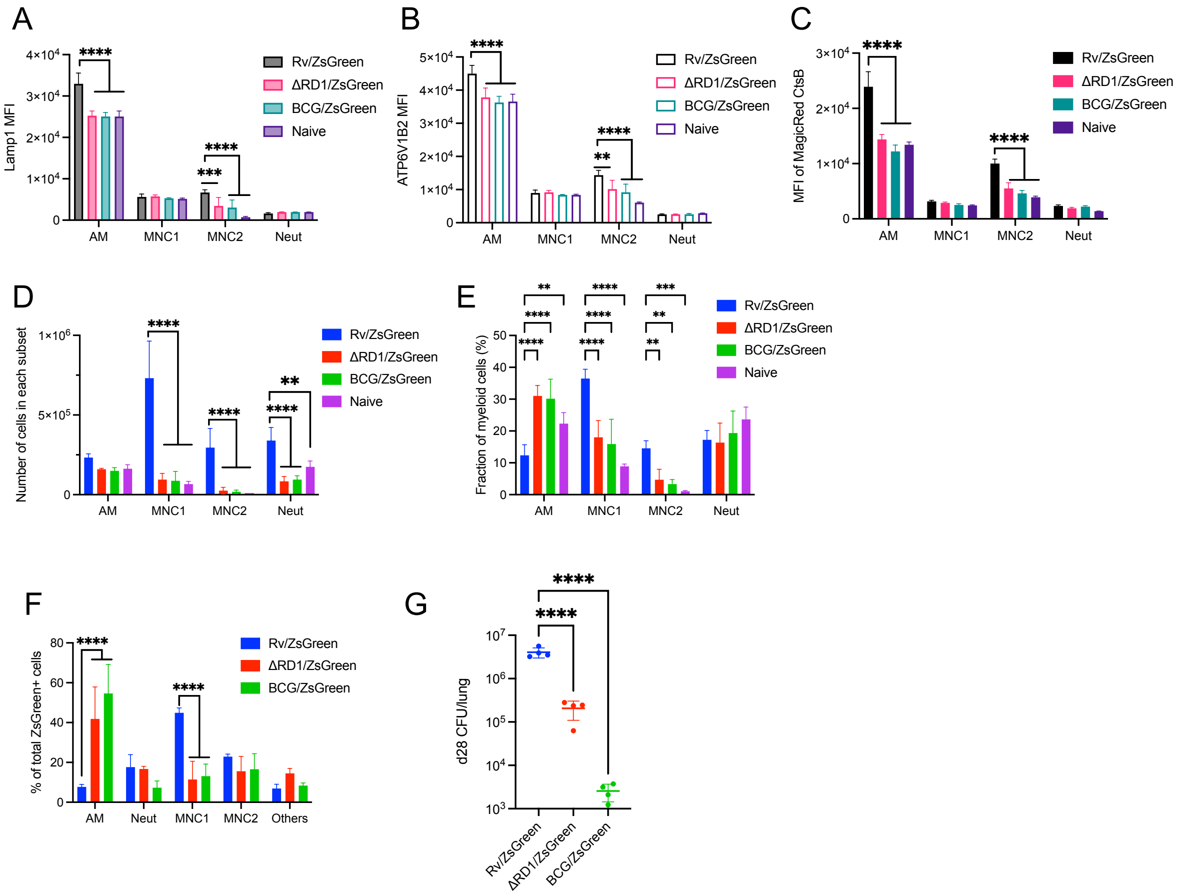


**Fig. S8. Mtb ESX-1 is required for MNC1 recruitment.**
C57BL/6 mice were infected with one of the three ZsGreen expressing strains via aerosol infection: Mtb H37Rv, Mtb H37Rv:ΔRD1, or *M. bovis* BCG. At 28 dpi, lungs were harvested for flow cytometry analysis or CFU assays. Naïve mice were uninfected.
(**A**) LAMP1 MFI of lung subsets from naïve mice and infected mice (28dpi).
(**B**) ATP6V1B2 MFI of lung subsets from naïve mice and infected mice (28dpi).
(**C**) MFI of fluorogenic CTSB product for lung subsets from naïve mice and infected mice (28dpi).
(**D**) Number of cells per subset from naïve mice, and mice infected with the indicated mycobacterial strains (28 dpi).
(**E**) Subset fractions of total myeloid cells (28 dpi).
(**F**) Frequency of cell types in total infected cells (28 dpi).
(**G**) Lung CFU for mice infected with different mycobacterial strains (28 dpi).
Results are presented as mean ± SD of 4-5 mice, representative of 2 independent experiments. **p<0.01 ****p<0.0001 by two-way ANOVA (A-F), or one-way ANOVA for (G).

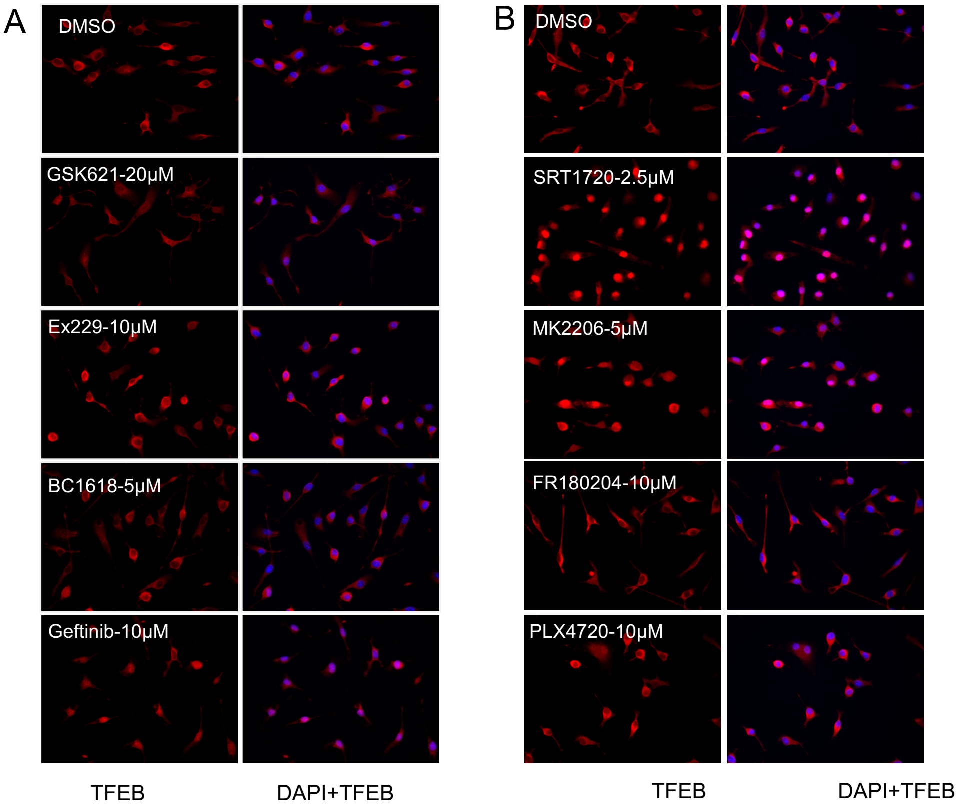


C

| Targets | Small molecules | Activation of TFEB nuclear translocation |
|---|---|---|
| c-Abl inhibitor | Imatinib | Yes |
| | Nilotinib | Yes |
| AMPK activator | GSK621 | Yes |
| | Ex229 | No |
| | BC1618 | No |
| EGFR inhibitor | Geftinib | Yes |
| Sirt1 activator | SRT1720 | Yes |
| AKT inhibitor | MK2206 | Yes |
| mTOR inhibitor | Torin1 | Yes |
| ERK inhibitor | FR180204 | No |
| Braf inhibitor | PLX-4720 | No |

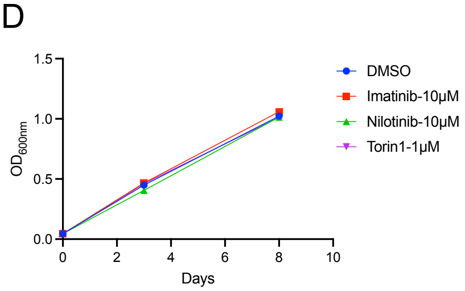


**Fig. S9. Effect of small molecules with distinct targets on TFEB activation.**

(**A-B**) BMDM were treated with indicated small molecules for 4h, then stained with DAPI and anti-TFEB for fluorescent microscopy.

(**C**) Summary of the effect of small molecules on the activation of TFEB nuclear translocation.

(**D**) Imatinib or Nilotinib do not inhibit H37Rv growth in 7H9 media. Three replicates per condition. Results are presented as mean ± SD, representative of 2 independent experiments.

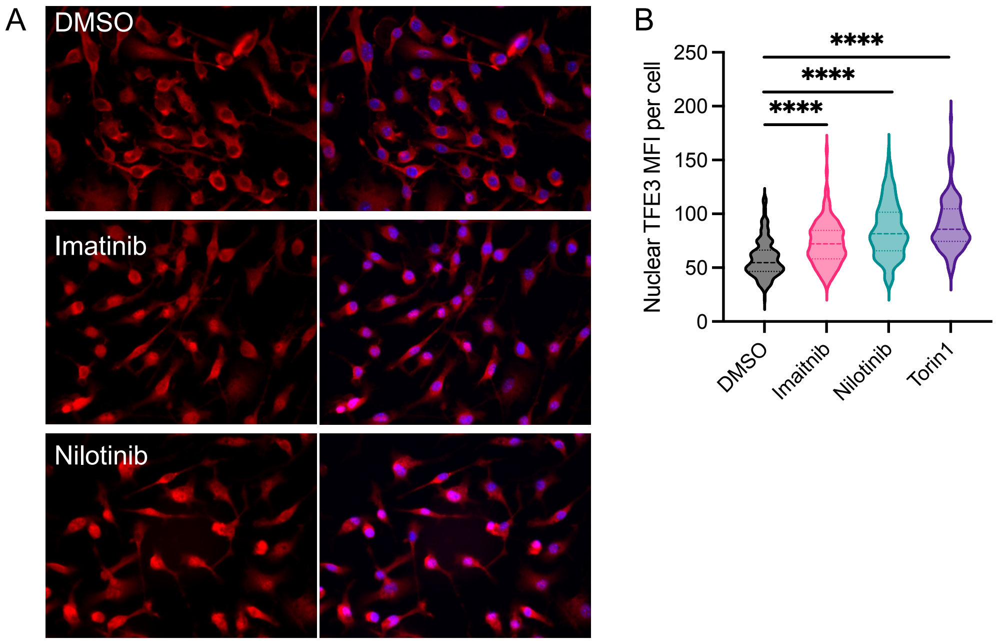


**Fig. S10. c-Abl inhibitors imatinib and nilotinib induce TFE3 nuclear translocation in BMDM.**
(**A**) BMDM were treated with indicated small molecules (Imatinib, 10 μM; Nilotinib, 10 μM) for 24h, then stained with DAPI and anti-TFE3 for fluorescent microscopy.
(**B**) Quantification of nuclear TFE3 MFI per cell from >127 cells for each condition in (A) using ImageJ.

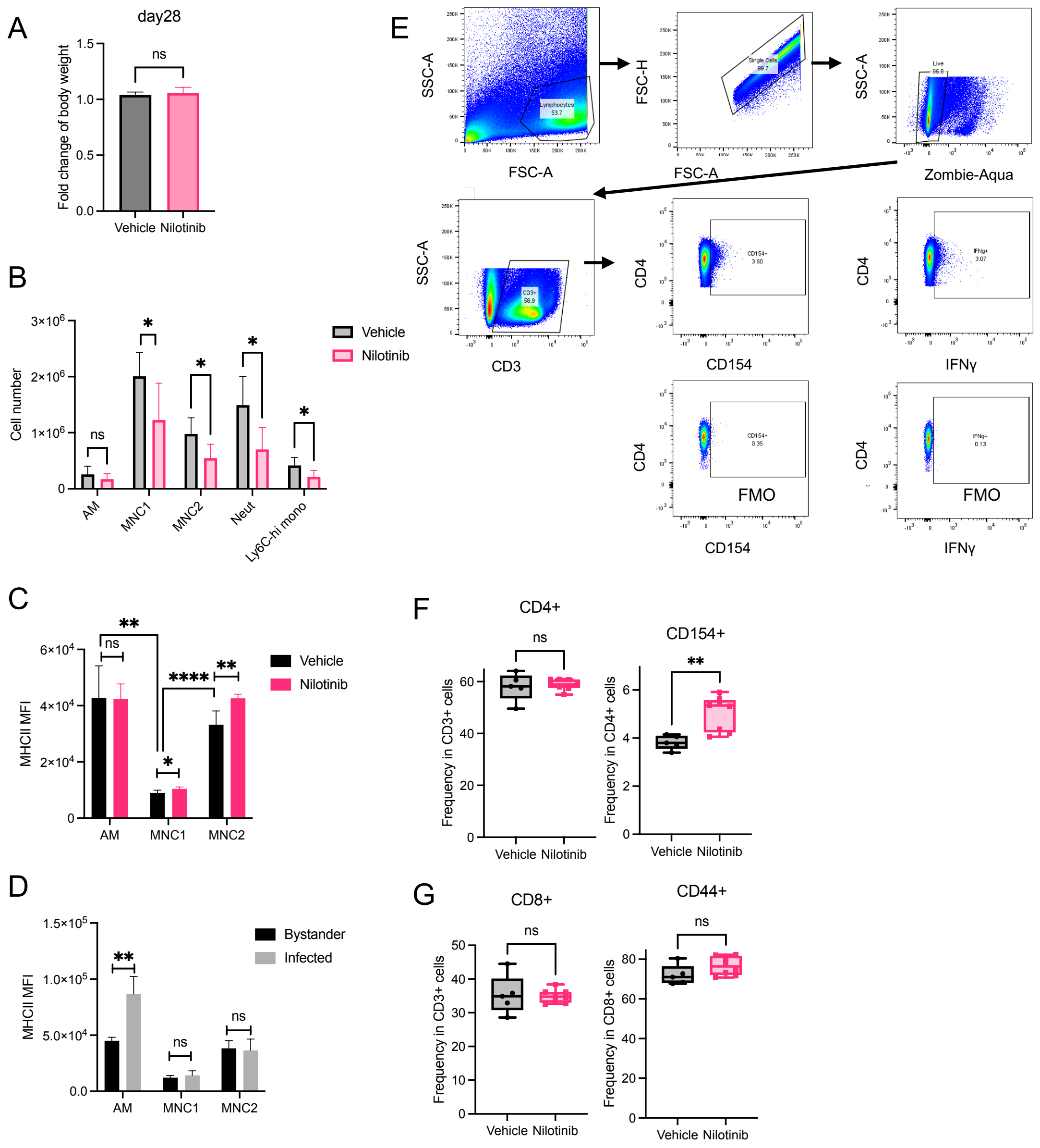


**Fig. S11. Effect of nilotinib treatment on mouse body weight, number of cells and MHCII level in lung myeloid subsets, and frequency of lung T cell subsets.**

Mice were infected with low-dose Mtb H37Rv-ZsGreen via aerosol, followed by treatment with vehicle or nilotinib (20mg/kg/day) intraperitoneally, 5 days/week, a total of 15 doses. Treatment was beginning on 7 dpi and ending on 27 dpi. Lungs were harvested for different assays at 28 dpi.

(**A**) Fold change of mouse body weight on 28 dpi relative to that on 7 dpi.

(**B**) Cell number in each lung subset from infected mice treated with vehicle or nilotinib (28 dpi).

(**C**) Surface MHCII level of different lung subsets from infected mice treated with vehicle or nilotinib (28 dpi).

(**D**) Surface MHCII level of infected and bystander cells for AM, MNC1, and MNC2 from infected mice without treatment (28 dpi).

(**E**) Gating strategy for defining lung $CD4^+$ T cells (28 dpi).
(**F**) Frequency of lung $CD4^+$ T cells, and frequency of $CD154^+$ cells in lung $CD4^+$ T cells (28 dpi).
(**G**) Frequency of lung $CD8^+$ T cells, and frequency of $CD44^+$ cells in lung $CD8^+$ T cells (28 dpi).
Results are presented as mean ± SD of 5-8 mice, representative of 2 independent experiments. *$p<0.05$, **$p<0.01$ by unpaired Student's t-test (A, F, G) or multiple unpaired Student's t-test (B-D).

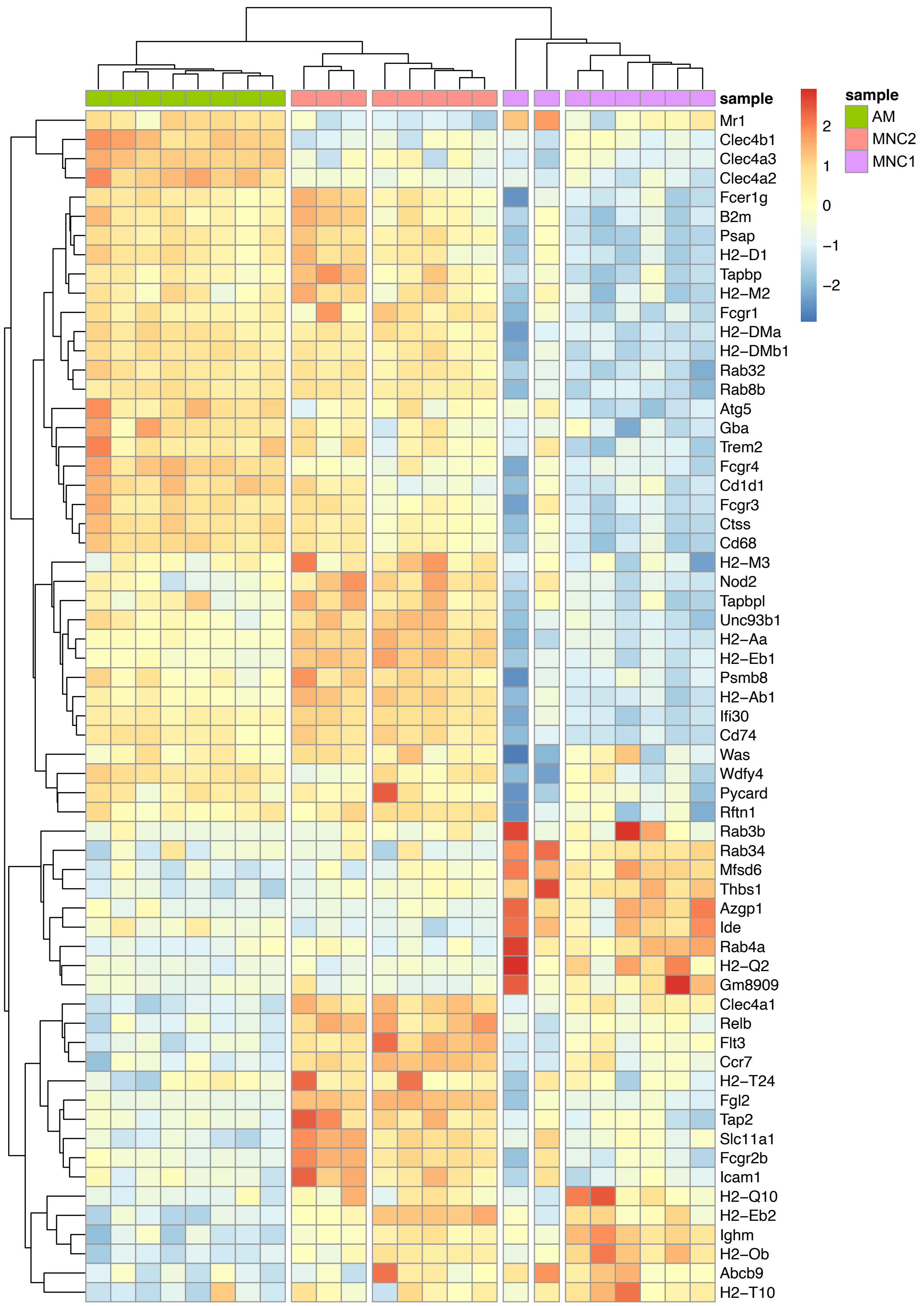


**Fig. S12. MNC1 are deficient in the expression of genes involved in antigen processing and presentation.** Heatmap was shown based on the Z-scores from variance stabilized read counts for significantly differentially expressed genes (adjusted p-value ≤ 0.05 and |log2 fold change|>0.5) of Gene Ontology term: antigen processing and presentation (GO:0019882).

**Table S1. Gene ontology analysis of differentially expressed genes (DEG) in MNC1 compared with those in other subsets (only antigen-related terms with an adjusted p-value <0.05 are shown)**

| Category | # of DEG in category | # of genes in category | term | p-value | Adjusted p-value | MNC1 vs |
|---|---|---|---|---|---|---|
| GO:0002478 | 24 | 32 | antigen processing and presentation of exogenous peptide antigen - BP | 4.59E-08 | 1.89E-04 | AM |
| GO:0002478 | 22 | 32 | antigen processing and presentation of exogenous peptide antigen - BP | 1.33E-08 | 5.90E-05 | MMC2 |
| GO:0002495 | 17 | 21 | antigen processing and presentation of peptide antigen via MHC class II - BP | 5.63E-07 | 0.00223852 | AM |
| GO:0002495 | 16 | 21 | antigen processing and presentation of peptide antigen via MHC class II - BP | 1.00E-07 | 0.00043502 | MMC2 |
| GO:0002501 | 12 | 14 | peptide antigen assembly with MHC protein complex - BP | 8.42E-06 | 0.0319096 | AM |
| GO:0002504 | 19 | 23 | antigen processing and presentation of peptide or polysaccharide antigen via MHC class II - BP | 9.15E-08 | 3.74E-04 | AM |
| GO:0002504 | 17 | 23 | antigen processing and presentation of peptide or polysaccharide antigen via MHC class II - BP | 1.21E-07 | 0.00052468 | MMC2 |
| GO:0019882 | 64 | 113 | antigen processing and presentation - BP | 1.32E-09 | 5.67E-06 | AM |
| GO:0019882 | 62 | 113 | antigen processing and presentation - BP | 1.09E-13 | 5.20E-10 | MMC2 |
| GO:0019884 | 27 | 39 | antigen processing and presentation of exogenous antigen - BP | 1.27E-07 | 5.16E-04 | AM |
| GO:0019884 | 27 | 39 | antigen processing and presentation of exogenous antigen - BP | 2.71E-10 | 1.24E-06 | MMC2 |
| GO:0019886 | 15 | 19 | antigen processing and presentation of exogenous peptide antigen via MHC class II - BP | 4.85E-06 | 0.01861506 | AM |
| GO:0019886 | 14 | 19 | antigen processing and presentation of exogenous peptide antigen via MHC class II - BP | 1.28E-06 | 0.005341 | MMC2 |
| GO:0048002 | 42 | 67 | antigen processing and presentation of peptide antigen - BP | 6.25E-09 | 2.64E-05 | AM |
| GO:0048002 | 38 | 67 | antigen processing and presentation of peptide antigen - BP | 8.04E-10 | 3.66E-06 | MMC2 |

**Table S2. Primers for qPCR analysis**

| Gene | Primer | Sequence (5'-3') | Product length/bp |
|---|---|---|---|
| Tfeb | Tfeb-F | TATCAGCTCCAACCCCGAGA | 147 |
| | Tfeb-R | CTGTACACGTTCAGGTGGCT | |
| Lamp1 | Lamp1-F | ACATCAGCCCAAATGACACA | 117 |
| | Lamp1-R | GGCTAGAGCTGGCATTCATC | |
| Ctsb | CtsB-F | TCCTTGATCCTTCTTTCTTGCC | 176 |
| | CtsB-R | ACAGTGCCACACAGCTTCTTC | |
| Ctsd | CtsD-F | AACGTGCTTCCGGTCTTTGA | 148 |
| | CtsD-R | GCTCCCCGTGGTAGTACTTG | |
| Hexa | Hexa-F | ACACCCTGTACCCCAACAAC | 154 |
| | Hexa-R | TGCTGTTTATTTGAGAAGCTGGG | |
| Atp6v0d2 | V0D2-F | CAGAGCTGTACTTCAATGTGGAC | 111 |
| | V0D2-R | AGGTCTCACACTGCACTAGGT | |
| Atp6v1b2 | V1B2-F | CTCTGGGGTGAATGGTCCAC | 137 |
| | V1B2-R | CCACTGCTTTGGAGCCACTA | |
| Gapdh | Gapdh-F | GGTGAAGGTCGGTGTGAACG | 87 |
| | Gapdh-R | GCAACAATCTCCACTTTGCCAC | |
